# Cross-species identification of cancer-resistance associated genes uncovers their relevance to human cancer risk

**DOI:** 10.1101/2021.05.19.444895

**Authors:** Nishanth Ulhas Nair, Kuoyuan Cheng, Lamis Naddaf, Elad Sharon, Lipika R. Pal, Padma S. Rajagopal, Irene Unterman, Kenneth Aldape, Sridhar Hannenhalli, Chi-Ping Day, Yuval Tabach, Eytan Ruppin

## Abstract

Cancer is an evolutionarily conserved disease that occurs in a wide variety of species. We applied a comparative genomics approach to systematically characterize the genes whose conservation levels significantly correlates positively (PC) or negatively (NC) with a broad spectrum of cancer-resistance estimates, computed across almost 200 vertebrate species. PC genes are enriched in pathways relevant to tumor suppression including cell cycle, DNA repair, and immune response, while NC genes are enriched with a host of metabolic pathways. The conservation levels of the PC and NC genes in a species serve to build the first genomics-based predictor of its cancer resistance score. We find that PC genes are less tolerant to loss of function (LoF) mutations, are enriched in cancer driver genes and are associated with germline mutations that increase human cancer risk. Furthermore, their expression levels are associated with lifetime cancer risk across human tissues. Finally, their knockout in mice results in increased cancer incidence. In sum, we find that many genes associated with cancer resistance across species are implicated in human cancers, pointing to several additional candidate genes that may have a functional role in human cancer.

## INTRODUCTION

Animal species are known to have dramatic differences in their cancer rates and lifespans, and several animals are considered cancer resistant while others are considered to be cancer prone (Gorbunova *et al*. 2014; Albuquerque *et al*. 2018). Studying the genomic underpinnings of these differences across various branches of life may provide insights into cancer development and cancer prevention/treatment options in humans (Seluanov *et al*. 2018).

The multistage carcinogenesis model states that “individual cells become cancerous after accumulating a specific number of mutational hits” (Seluanov *et al*. 2018; Nordling, 1953). Based on this model, larger (and longer-living) animals are expected to have higher cancer incidence as they have more stem cell divisions overall, resulting in a higher likelihood of producing and propagating carcinogenic mutations. For humans, it has been shown that the risks of cancer development across different tissue types are correlated with their corresponding estimated number of lifetime stem cell divisions (Tomasetti *et al*. 2015 and 2017); consistent with that, human cancer risk is indeed correlated with body height (Khankari *et al*. 2016). However, cancer risk does not correlate with body size across species, a contradiction known as Peto’s paradox (Peto, 1947; Tollis *et al*., 2017; Seluanov *et al*. 2018). For example, humans do not have higher cancer risk than mice despite having thousands of times more cells (Lipman *et al*. 2004; Szymanska *et al*. 2014; Ikeno *et al*. 2009). More drastically, the cancer-resistant bowhead whale (Keane *et al*., 2016) can weigh 100 tons, live for over 200 years (George *et al*., 1999) and have millions times more cells than mice. It follows that different species must have evolved different cancer resistance mechanisms to fit their lifestyles, modifying the “baseline” probability of malignant transformation determined by body size, lifespan, and tissue stem cell division (see Supp. Note for a short review of such mechanisms).

Numerous studies have adopted comparative genomics approaches to understand the evolution of cancer resistance mechanisms across mammals. Some have focused on known human cancer genes and their homologs. For example, Vicens and Posada (2018) found that genes related to DNA repair and T cell proliferation have evolved under positive selection in mammals. Tollis *et al*. (2020) found that the number of paralogs of human cancer genes across mammals is positively correlated with the species’ lifespan, but not body size. Vazquez and Lynch (2021) reported wide-spread tumor suppressor gene (TSG) duplications across both large and small Afrotherian species. Other studies focused on body size and longevity, yielding some insights into Peto’s paradox. Kowalczyk *et al*. (2020) analyzed genes whose evolutionary rates across mammals correlate with body size and lifespan and discovered cancer resistance-related genes that are under increased evolutionary constraints in larger and longer-living mammals. Ferris *et al*. (2018) identified regions with accelerated evolution in specific mammals, including several cancer resistant species, which provided some insights on the cancer resistance mechanisms they have developed.

Unlike previous studies that focused exclusively on mammals, here we perform a comprehensive genome-wide comparative study aimed at identifying genes related to cancer resistance across a wide range of vertebrate species. To this end, we estimated the protein conservation scores across species including mammals, birds and fish, identifying genes whose conservation levels are associated with cancer resistance estimates. We then use these cancer-resistance associated genes to build the first genomics-based predictor of cancer resistance for any species. We show that the biological processes associated with cancer resistance vary across taxonomic groups (classes and orders of species), pointing to the diversity in the evolutionary paths and mechanisms for resisting cancer. Finally, the genes identified from this phylogenetic analysis are enriched for cancer driver genes and in genes associated with cancer risk in humans. These results show that a comparative genomic approach can help identify genes involved in human cancers.

## RESULTS

### Computing *gene conservation* and *species cancer-resistance* estimates

We computed a matrix (Tabach *et al*. Nature 2013; Tabach *et al*. MSB 2013) of gene conservation scores (phylogenetic profiles) across 240 species for which we had phenotypic information in the AnAge database (Tacutu *et al*. 2018) and sequence information from UniProt (UnitProt Consortium, 2021), Refseq (O’Leary *et al*. 2016), Keane *et al*. (2015), and NCBI (Sayers *et al*. 2021) databases. To do this, the protein sequence similarity between each gene in the genome of a reference species and its orthologs in each of the rest of the species (termed phylogenetic profiling; Pellegrini *et al*. 1999) was measured using the bit score computed with BLASTP (Altschul *et al*. 1990). The BLASTP bit scores were normalized by their gene length (Tabach *et al*. Nature 2013; Sherill-Rofe *et al*. 2019) and then rank-normalized across all genes within each species to control for the evolutionary distance between the reference and each species (Methods). These rank-normalized values range from 0 to 1, with higher values corresponding to higher conservation levels. This method is termed rank-based phylogenetic profiling. We primarily focused on the human as the reference species (Braun *et al*., 2020) as we are interested in making our findings relevant to human cancers. However, we demonstrated that our conclusions are robust to the choice of reference (Methods, Supp. Note), largely because the normalization effectively removes dependency on phylogenetic distance.

Since the cancer incidence rates of most species are largely unknown, we used two proxy cancer-resistance estimates that have been proposed in the literature – *MTLAW* and *MLCAW*. MLTAW assumes that the level of cancer resistance in a given species needs to roughly counteract its risk of cancer development due to cell division, which is proportional to ML^6^ × AW, where ML denotes the species maximum longevity and AW denotes its adult weight (Peto *et al*. 1977, 2015; Vazquez *et al*. 2021; Methods). MLCAW considers the well-established correlation between lifespan and body weight (AW) across many species (Speakman, 2005) and thus regresses out the species AW from its ML (Methods). We computed MLTAW and MLCAW for 193 out of the 240 species for which both ML and AW data was publicly available (Table S1, Methods). These 193 species are from multiple Vertebrata classes, including Mammalia (mammals, n=108), Aves (birds, n=55), Teleostei (teleost fishes, n=18), and Reptilia (reptiles, n=7).

### Genes associated with cancer resistance are enriched in cell cycle, DNA repair, immune response, and different metabolic pathways

For each gene, we computed the Pearson correlation coefficient between its conservation scores and the cancer-resistance estimates (MLTAW and MLCAW) across all species (Tables S2A,B; Methods). We then computed the pathway enrichment of the positive and of the negatively correlated genes (termed PC or NC genes, respectively) (Tables S3A,B; Methods). PC genes correlated with either the MLCAW (**Fig. 1**) and MLTAW measures (Fig. S1) are enriched for cell cycle, immune response, DNA repair, and transcription regulation pathways (FDR<0.1), indicating that many genes in these pathways are more conserved in the relatively long-lived cancer-resistant species. NC genes are enriched for a diverse range of metabolic pathways (FDR<0.1, **Figs. 1**,S1).

**Figure 1:**
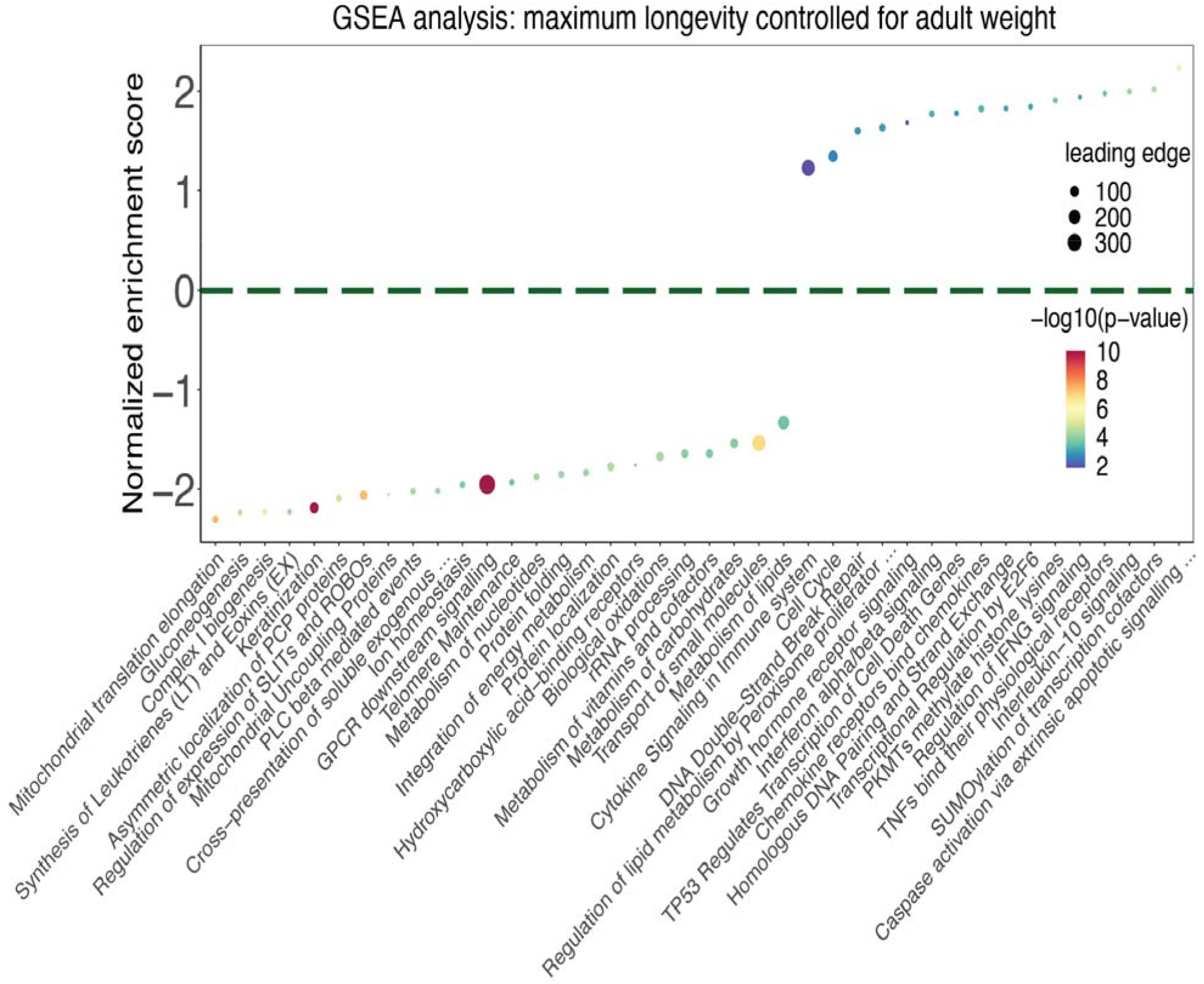
Summary of the top significantly enriched pathways (adjusted P<0.1) by the genes whose conservation scores are correlated with cancer-resistance estimates, using gene set enrichment analysis (GSEA) with gene set annotations from the Reactome database. The cancer-resistance estimate used is “Maximum longevity controlled for adult weight” (MLCAW). Normalized enrichment score is plotted on the Y-axis, where positive values correspond to enrichment by the positively correlated (PC) genes and negative values correspond to enrichment by the negatively correlated (NC) genes. The dot color represents the significance of the enrichment (negative log10 GSEA P value), and the dot size represents the number of genes in the “leading edge”, i.e. the set of genes that are enriched in a pathway. For the sake of clarity, only a subset of the enriched pathways (FDR<0.1) are shown and long pathway names have been shortened (using “…”). The complete pathway enrichment results are given in Table S3B.

### PC and NC gene conservation scores are predictive of species cancer resistance

We next asked whether it is possible to accurately predict the cancer-resistance estimates of individual species from their gene conservation scores. For a species, given the median conservation score (MCS) of all its genes, we defined a *cancer resistance (CR) score* that quantifies how many of the PC genes have conservation scores > MCS and how many NC genes have conservation scores < MCS (normalized by the total number of genes, Methods). Using a standard leave-one-out cross-validation (LOOCV) procedure, both across all species and then focusing on mammals or birds (as these groups contain sufficient number of species), we find that the CR score is strongly predictive of the cancer-resistance estimates of a left-out species using the PC/NC genes identified from the other species (all species: MLTAW Spearman’s ρ=0.44, P=1.32e-10, Fig. S2, MLCAW ρ=0.51, P=2.31e-14, **Fig. 2A;** mammals: MLCAW ρ=0.67, P=1.58e-15, **Fig. 2B,** MLTAW ρ=0.76, P=8.99e-22, **Fig. 2C;** the results for birds are provided in Supp. Note and Fig. S3). Supp. Note and Figs. S4-S8 present both technical controls (choosing random sets of PC and NC genes to predict cancer-resistance estimates), and robustness analysis showing that these results hold when (i) using two-fold cross-validation instead of LOOCV, (ii) under changes in the choice of reference species and threshold parameters, (iii) using alternate predictors showing the contributions of PC or NC genes separately and finally (iv) using Spearman’s instead of Pearson correlation to identify PC/NC genes; Tables S1, S2; Methods). The predicted CR scores learnt from all mammals (LOOCV) also show significant correlation within different subgroups (as an example, MLCAW Spearman’s ρ=0.85, P=0.0061 for the order Chiroptera, i.e. bats, **Figs. 2D;** for others see Supp. Note, Fig. S9). Similarly, the predicted CR scores learnt from all birds’ species (LOOCV) show significant correlation within the order Passeriformes for which we have the largest number of samples (Spearman’s ρ=0.79, P=0.0012, Fig. S3B, Supp. Note).

**Figure 2:**
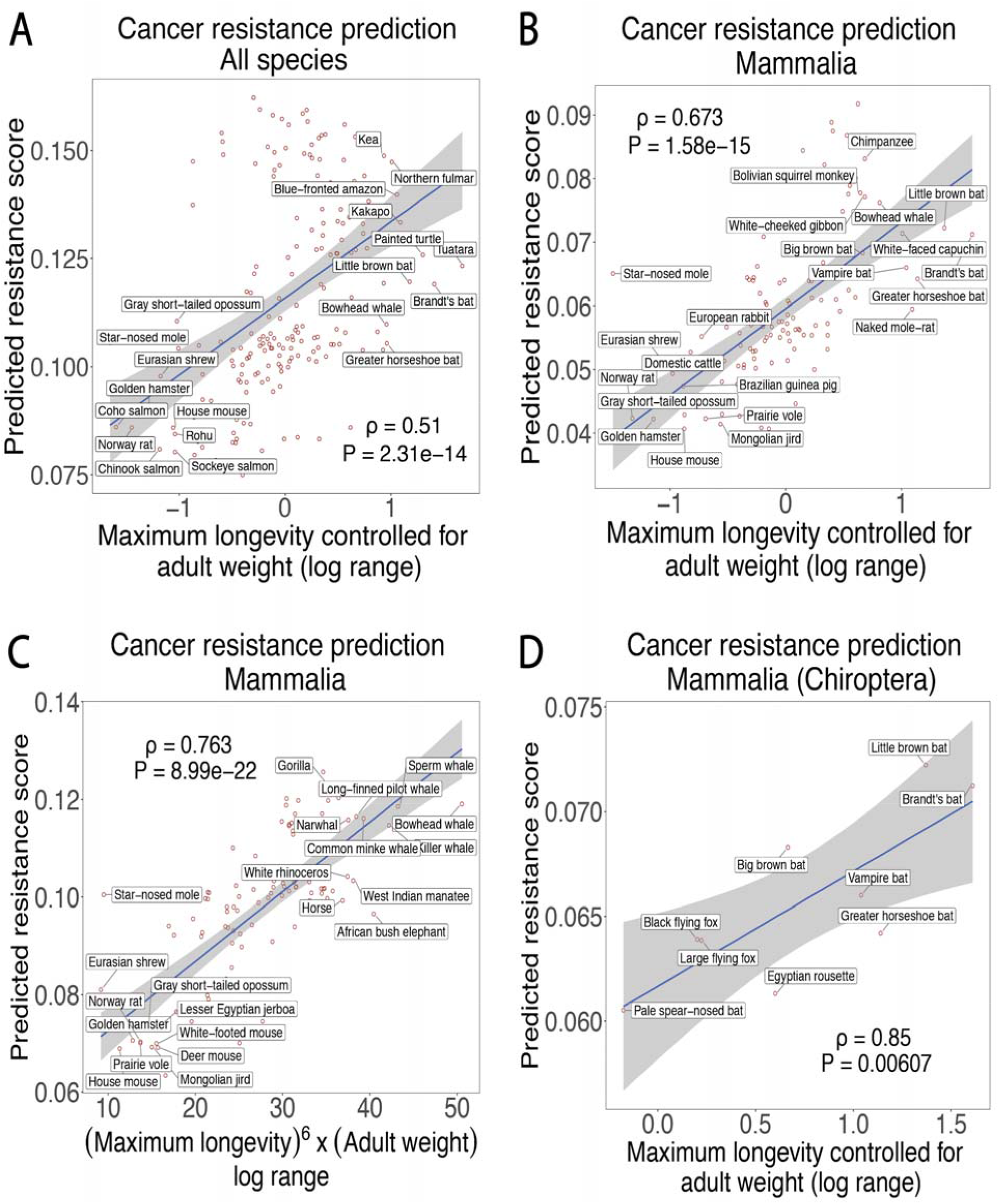
Scatter plots showing the correlation between the predicted cancer resistance (CR) scores computed based on gene conservation (Y-axes) and either of the two cancer-resistance estimates (X-axes): MLCAW i.e. maximum longevity controlled for adult weight) or MLTAW, i.e. (Maximum longevity)^6^ x (adult weight), with leave-one-out cross-validation. Results for (A) MLCAW across all species; (B) and (C) MLCAW and MLTAW within mammalian species, respectively; (D) using the MLCAW mammalian-specific predictions only within a subgroup: order Chiroptera. Species with the top and bottom 5% MLCAW values in (A), the top and bottom 10% MLTAW or MLCAW values in (B,C), all data-points in (D), are labeled by their common names. In each panel, the Spearman’s ρ and p-values (P) are shown.

Our results show that high CR scores are predicted for many long living species that are considered to be cancer resistant, including the bowhead whale, the African elephant, the chimpanzee, the Brandt’s bat, the naked mole rat, etc **(Figs. 2A-D,** S9, and S6; Supp. Notes; Gorbunova *et al*. 2014; Varki & Varki, 2015; Seluonov *et al*. 2018; Wilkinson & Adams, 2019). Predictions of cancer resistance in additional species without documented body weight or life span are provided in Table S1. The PC/NC genes derived from one clade do not however yield accurate predictions in another taxonomic group (across classes: Fig. S10; across mammalian orders: Fig. S11; Tables S2, S4). This indicates that different taxonomic groups may have evolved to have some differences in their cancer resistance mechanisms, which we study next.

### Cancer resistance-associated genes in mammals, birds, and teleost fishes

We next repeated the correlation analysis between gene conservation score and MLTAW/MLCAW scores separately for mammals, birds, and the teleost fish, and computed the PC/NC gene-enriched pathways for each of the three groups (Methods). There are overall significant overlaps among the NC gene enrichments of the three classes, especially based on MLCAW (odds ratio, i.e. OR as large as 18.9, Fisher’s exact test adjusted P as small as 1.8e-11; Figs. S12A,B; Table S3I), while the overlap among the PC gene sets are mostly insignificant (other than between mammals and birds using MLCAW: OR=5.06, adjusted P=0.037; Figs. S12A,B; Table S3I). Both common pathways (e.g. GPCR signaling) and pathways unique to specific classes (e.g. fatty acid and amino acid metabolism, PI3K-AKT signaling pathway in birds) were observed (details in **Figs. 3A,** S12C, Table S3).

**Figure 3:**
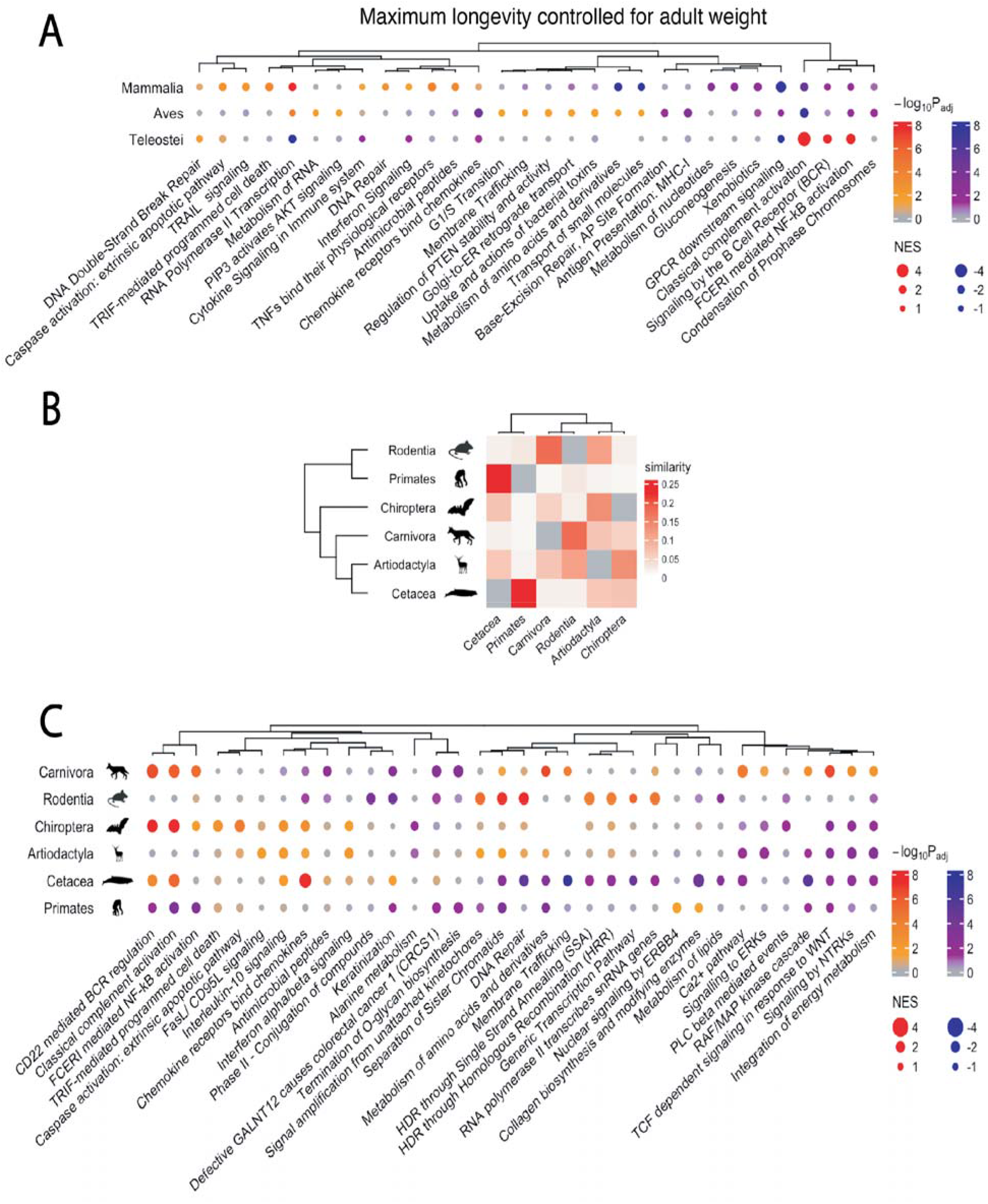
Gene set enrichment analysis (GSEA) of gene conservation correlations with the cancer-resistance estimate “maximum longevity controlled for adult weight” (MLCAW) specifically in different taxonomic groups. **(A)** A summary visualization of the top enriched pathways (with GSEA) based on gene conservation correlations with MLCAW in Mammalia (mammals), Aves (birds) and Teleostei (teleost fishes). A selected subset of top gene sets are shown to save space, all with adjusted P<0.1 in at least one of the classes (Methods). GSEA significance (negative log10; adjusted P values) is encoded by dot color, with two sets of colors (red-orange and blue-purple) representing positive or negative enrichment, respectively; grey color means adjusted P>=0.1. Dot size represents the absolute value of normalized enrichment scores (NES) measuring the effect size of enrichment. The complete GSEA results are given in Table S3. **(B)** A heatmap showing the similarity (Jaccard index) between the significantly enriched gene sets (FDR<0.1) from each pair of mammalian orders, based on the MLCAW correlation. The dendrogram on the left is the phylogenetic tree of the mammalian orders, and the rows of the heatmap are arranged accordingly. The dendrogram on the top represents the hierarchical clustering of the orders based on their similarities in the GSEA results. **(C)** A summary visualization of the top enriched pathways (with GSEA) based on gene conservation correlations with MLCAW in different mammalian orders. This figure panel should be read as in **(A)**.

The class Mammalia contains the largest number of species (n=108) with available data, allowing us to further investigate the specificities in several orders, including Rodentia (rodents, n=20), Primates (n=18), Carnivora (carnivores, n=18), Artiodactyla (even-toed hoofed mammals, n=11), Cetacea (aquatic mammals like whales, n=10), and Chiroptera (bats, n=9). **Figs. 3B** and S12D visualize the similarities (using a Jaccard index-like measure) between the significant PC/NC gene-enriched pathways from pairs of orders (Methods). The different orders exhibit an overall similarity pattern that does not fully coincide with their phylogenetic relations (dendrograms in **Figs. 3B,** S12D). Primates share the highest pathway-level similarity with Cetacea (Fisher’s exact test adjusted P<2.2e-16; Table S5). Rodentia appears the most similar to Carnivora (Fisher’s exact test adjusted P<2.2e-16; Table S5) and Artiodactyla. However, specific enriched pathways are shared across orders (Table S5; **Figs. 3C,** S12E). This includes various cytokine signaling pathways and extrinsic apoptotic pathways that are mostly enriched by PC genes **(Figs. 3C,** S12E), recapitulating the role of the innate immune system in the evolution of more cancer-resistant mammalian species. WNT and VEGF signaling, and lipid metabolism are among the pathways showing consistent NC gene-enrichment across orders (esp. based on MLTAW, **Figs. 3C,** S12E). Interestingly, DNA repair-related pathways, showing PC-enrichment in Rodentia and other orders, exhibit very strong NC-enrichment in Cetacea (based on MLCAW, **Fig. 3C).** Complement cascade/activation also exhibit an order-specificity **(Figs. 3C,** S12E). These observations point to the diversity in pathways associated with cancer resistance in different mammalian orders.

### Cancer resistance-associated genes are enriched for human cancer driver genes

We turned to ask whether PC and NC genes are enriched for well-established human cancer driver genes (from the COSMIC database, Forbes *et al*. 2015). PC genes (but not NC genes) inferred either across all species or mammals are highly enriched for human tumor suppressor genes (TSGs; adjusted P=0.0011 and 0.013, respectively; **Fig. 4A;** Table S6B), and oncogenes in the all-species analysis (adjusted P=0.0011, **Fig. 4A).** These strong enrichments still hold with PC genes identified while excluding all primates (Table S6B). We note that excluding the human TSGs and oncogenes from the PC/NC genes when computing the CR score does not reduce the accuracy in predicting cancer resistance across species **(Figs. 4B** and S9). Finally, we find that the PC genes inferred across all species are enriched for the genes reported in various human cancer GWAS studies curated from the EBI GWAS Catalog (GSEA adjusted P=0.02; enrichments still hold with PC genes identified after excluding all primates; Table S6C; results were obtained using the *MLCAW* measure, Methods).

**Figure 4.**
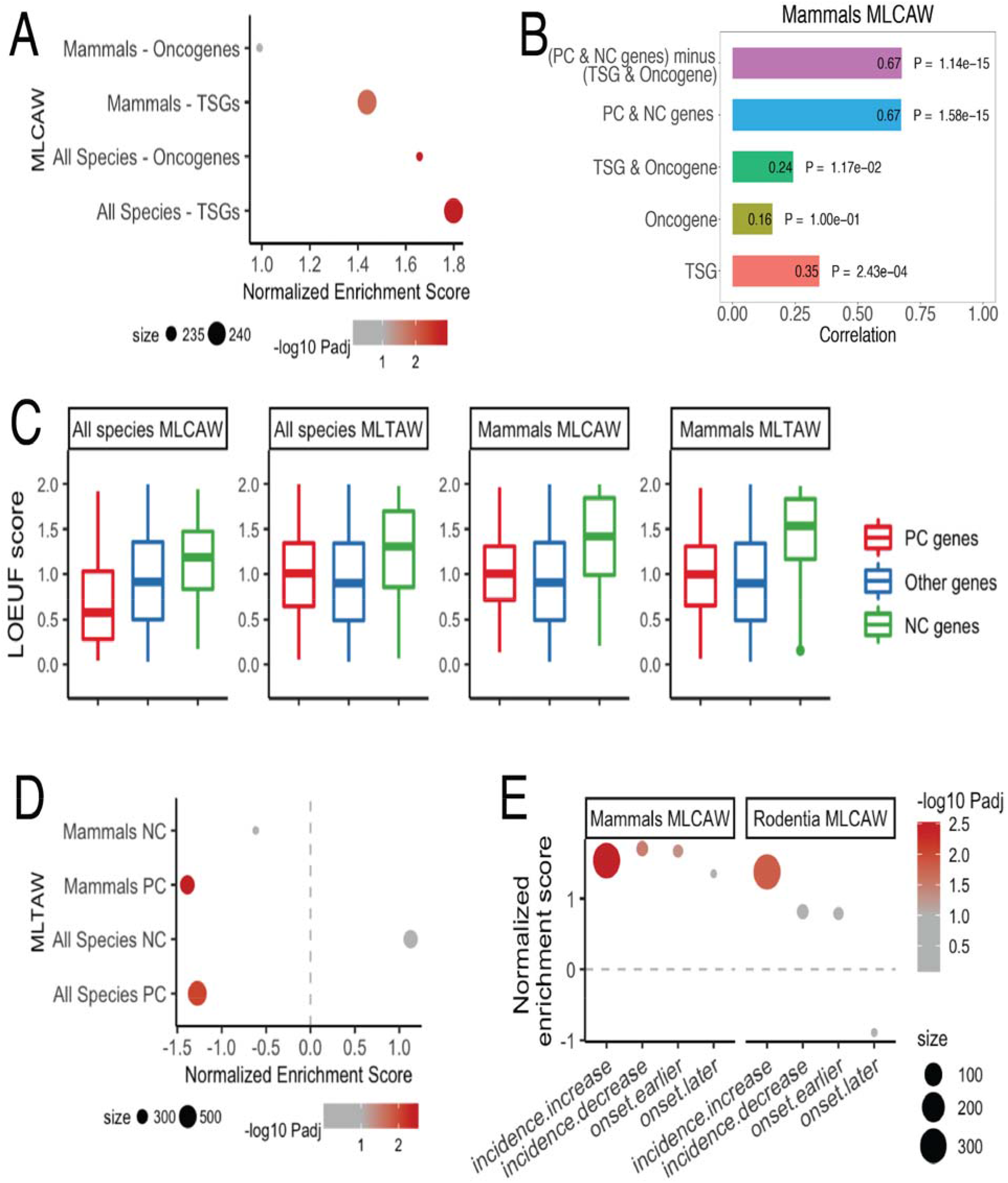
**(A)** A summary of enrichment via gene set enrichment analysis (GSEA) results for human tumor suppressor genes (TSGs) or oncogenes whose conservation scores correlate with MLCAW measure in all species or in mammals. Dot size corresponds to gene set size. Dot color denotes negative log10 adjusted P value from GSEA; grey corresponds to adjusted P>=0.1. Positive normalized enrichment score (Y-axis) corresponds to enrichment by PC genes, and vice versa for NC genes. **(B)** Spearman’s correlation (ρ) in predicting cancer resistance (MLCAW) in mammals using only TSGs, only oncogenes, both TSGs and oncogenes, using PC and NC genes in cross validation, using PC and NC genes after removing TSGs and oncogenes in cross validation is shown (Methods). **(C)** Box plots comparing the LOEUF scores of the genes whose conservation score positively (PC) or negatively correlates (NC) with a cancer-resistance estimate, and the other genes in the genome, based on the two cancer-resistance estimates (maximum longevity)^6^ x (adult weight) (MLTAW), and the residue of maximum longevity after regressing out adult weight (MLCAW), either in all species or in mammalian species. **(D)** A summary of the GSEA results on the enrichment of the top PC/NC genes from the MLTAW correlation in all species or mammals for genes whose expression levels correlate with the tissue-specific cancer incidence across human tissues (SEER, 2018). Dot size and color are interpreted as in (A). Positive normalized enrichment score (X-axis) corresponds to enrichment by genes whose higher expression is associated with higher cancer incidence across human tissues, vice versa. **(E)** A summary of enrichment (via GSEA) for mice genes whose knockout cause cancer-related phenotypes in the genes whose conservation scores correlate with MLCAW in all mammals or specifically rodents. “incidence.increase” denotes the mice genes whose knockout results in an increase in observed cancer incidence obtained from the MGI database, similarly for other gene sets listed on the X-axis. Dot size, color, and the normalized enrichment score (Y-axis) are interpreted as in (A).

To study the nature of selection operating on the PC and NC genes in human evolution, we compared the LOEUF (loss-of-function observed/expected upper bound fractions) scores of the PC, NC, and the rest of the genes (background) in the human genome; the higher the LOEUF score, the greater the tolerance to loss-of-function mutations (Karczewski *et al*. 2020). We find that the NC genes have significantly higher LOEUF scores compared to PC genes and the rest of the genes in the genome **(Fig. 4C;** Table S6D), indicating that they are subject to weaker purifying selection pressure than the PC and other genes, as expected.

### The expression of PC genes in normal human tissues is associated with their lifetime cancer risk

As PC genes are enriched for human TSGs and oncogenes, they may also have roles in modulating human cancer risk. We hence examined whether their expression levels across different non-cancerous human tissues are associated with lifetime cancer risks across these tissues, which are highly variable (Tomasetti & Vogelstein, 2015). Analyzing lifetime risk data (the SEER program, 2018 and Tomasetti & Vogelstein, 2015) and the GTEx RNA-seq data (Lonsdale *et al*. 2013) we find that the *MLTAW* PC genes (but not *MLCAW* ones) are enriched for genes whose expression levels negatively correlate with cancer risk across tissues (adjusted P=0.0088 in the all-species analysis and 0.003 in the mammal-specific analysis, **Fig. 4D;** results still hold after excluding primates when identifying the PC genes; Table S6E). We do not see a similar pattern using NC genes **(Fig. 4D).**

### PC genes are associated with cancer incidence in mice and canine transmissible venereal tumors

We investigated the relevance of PC and NC genes to cancer risk in other mammalian species. We first focused on the mouse, which has been extensively studied genetically. Mining the MGI database (Bult *et al*. 2019), we assembled lists of genes whose knockout in the mouse results in cancer-related phenotypes including the increase/decrease of cancer incidence and cancer onset time (Methods). We find strong enrichment of the MLCAW PC genes (in all mammals, and specifically rodents) in cancer incidence-increasing genes (P=0.003, **Fig. 4E;** Table S6F). In the all-mammal analysis, however, a weaker PC-enrichment was observed for incidence-decreasing genes and “earlier onset” genes (adjusted P<0.05, **Fig. 4E).**

To investigate the role of PC genes in tumorigenesis, we analyzed the expressed mutated genes in a single-cell phylogeny of a mouse melanoma model (Pérez-Guijarro *et al*., 2020), in which five subclones (B1 to B5) were identified (Mehrabadi *et al*., 2021). The mutated genes are significantly enriched with the PC genes from the all-species MLTAW and MLCAW analysis (Table S7), consistent with the putative function of PC genes as safeguards of cellular transformation. Interestingly, the mutated PC genes in each subclone are enriched in distinct pathways (Table S7), implying that, following the initial common mutations, each subclone evolved independently by overcoming different cancer resistant mechanisms. These results illustrated how PC genes are involved in the carcinogenic process.

Additionally, we investigated canine transmissible venereal tumors (CTVTs), a naturally occurring transmissible cancer in dogs that first arose about 11,000 years ago (Murchison *et al*. 2014). In CTVTs, more than 10,000 genes carry nonsynonymous mutations, and 646 genes have LoF via different mechanisms (Murchison *et al*. 2014). Notably, there is a significant enrichment of the PC genes from the mammals MLTAW analysis for CTVT LoF genes (adjusted P=0.017, Fig. S14; Table S6G).

### Specific PC genes with strong evidence of cancer-relevance across many different analyses

We manually curated the lists of PC genes, identifying a subset showing relevance to cancers based on multiple criteria according to the various analyses performed above (e.g., being human cancer drivers, genes whose knockout results in cancer-related phenotypes in mice, etc. Methods; Table S8), and investigated their functions closely. Several of these curated genes have known or investigated associations to germline cancer risk syndromes. For instance, mutations in *BRCA1* and *BRCA2* are extremely well-established in defining hereditary breast and ovarian cancer syndrome (Chen & Parmigiani, 2008; Kuchenbaecker *et al*., 2017). Risk syndromes have been defined more recently for moderate penetrance genes such as *CHEK2* (breast and colon cancer) and *BRIP1* (ovarian cancer) (Cybulski *et al*. 2011, Ramus *et al*. 2015). *NBN* is currently under investigation for contribution to germline breast and ovarian risk (Zhang *et al*., 2011, Kurian *et al*., 2017). Some of the manually prioritized genes we identified are currently under investigation for association to cancer risk, and our results may support greater consideration of their contribution to human cancer development. For example, *BUB1B*, prioritized strongly in our list, is under investigation for association with early-onset colorectal cancer (Hahn *et al*. 2016) but does not have clinically relevant screening or management recommendations at this time.

Other curated genes have known clinical associations to cancer. *NPM1* and *TET2* are currently used for prognostication with acute myeloid leukemias (Verhaak *et al*. 2005; Chou *et al*. 2011). BCG, a therapy used in early-stage bladder cancer, is a ligand for TLR2 (Fuge *et al*. 2015; Urban-Wojciuk *et al*., 2019). Interferon gamma (IFNG) is currently being evaluated therapeutically with other immunotherapies across multiple trials (Ni *et al*. 2018), and mutations in *DEK* are currently being used as biomarkers in multiple hematologic trials (Sanden & Gullburg, 2015). Numerous genes in our curated list (Table S8), while linked to cancer as per our enrichment analysis, have not yet had their functional relevance clarified, such as *RBM27, STAM2, SCAF4, SP140, RSBN1, SECISBP2L, THUMPD2, PIFO* and *POLK*. These genes may warrant higher prioritization to study their role across human cancers and potential therapeutic relevance based on our findings.

## DISCUSSION

We systematically analyzed the genomes of almost 200 species to identify genes whose conservation levels are correlated with cancer resistance estimates across different taxonomic groups and characterized their functional enrichment. We built the first genomics-based predictor of cancer resistance across species. We further studied the relevance of these phylogenetically derived cancer resistance-associated PC/NC genes to cancer development in humans.

Overall we found that PC genes are highly relevant to carcinogenesis and enriched with cell cycle, DNA repair, immune response and transcription regulation genes in the all-species analysis (Fig. 1 and S1). These results echo those of a recent study showing that cell cycle, DNA repair, NF-κB-related, and immunity pathways have higher evolutionary constraints in larger and longer-living mammals (Kowalczyk *et al*. 2020). MLTAW and MLCAW were used as two species cancer-resistance estimates, and, per definition they are correlated with each other (Spearman’s ρ=0.45, P=4.12e-11). However, despite the overall similarity at a high level, the MLTAW and MLCAW analyses uncover different aspects of the cancer resistance mechanisms. The top PC-enriched pathways using the MLTAW measure, where both body size and lifespan are multiplication factors, are dominated by cell cycle regulation and transcription/RNA regulation (Fig. S1), suggesting a stronger role of tissue stem cell division. The MLCAW measure, however, controls for body size, and its PC-enriched pathways include innate immunity or cell death for eradicating defect cells (Woo *et al*. 2015), highlighting the involvement of these factors after reaching adult size. NC genes computed with both MLTAW and MLCAW are notably enriched for processes related to cell metabolism, indicating either evolutionary metabolic constraints in the smaller/shorter-lived species or accelerated evolution of metabolism in the larger/longer-lived species (Speakman 2005).

Another notable pattern is the variability in the PC/NC gene functions across different taxonomic groups -- it’s frequently observed that genes of one pathway can be PC in one group but NC in another. Such variation may reflect trade-off between individual lifespan and survival/reproduction function dependent on the different lifestyles in different groups of species. Some of the observed order-specific enrichments are consistent with known mechanisms of cancer resistance for the corresponding species. For example, the naked mole rat is known to have more efficient excision repair systems than the mouse (Evdokimov *et al*. 2018) and an active complement system has been observed in bats (Moore *et al*. 2011; Mellors *et al*. 2020).

We outline several limitations of our study. First, the gene conservation computation is based on comparison to a reference species and rank normalization, which does not consider paralogous genes or the phylogenetic tree structure. While alternative methods may be used to adjust for the inter-phylogeny distances, we showed that our results are robust to the choice of the reference species (e.g., with house mouse and thirteen-lined ground squirrel as references; mouse is a known cancer-prone species, unlike human; Supp. Note) and various other conservation scoring parameters. Additionally, most of our downstream analyses were on the pathway-level, which mitigates the potential variation due to paralogs. Second, while we have explained the rationale for MLTAW and MLCAW as proxy species cancer-resistance estimates, these estimates cannot capture variations in cancer resistance that are not reflected through body size and lifespan, e.g., those related to adaptation to different oxygen and oxidative stress levels (Supp. Note; Hammarlund *et al*. 2018). Lastly, although the knockout mouse data validates the cancer-resistance function of many PC genes (Fig. 4E), further studies are obviously required for testing the roles of PC and NC genes (and our curated gene list in Table S8) in human carcinogenesis.

In summary, this study presents a systematic species comparison identifying key genes and pathways associated with cancer resistance across species. Many of the genes identified are implicated in human cancers, and their further study may increase our understanding of human cancer development, prevention and treatment.

## METHODS

### Computation of gene conservation scores

We created a matrix of conservation scores for 20076 genes across 240 species with human genome as a reference. The amino acid sequence of the proteins in all of these species are available in UniProt (UnitProt Consortium, 2021), Refseq (O’Leary *et al*. 2016), Keane *et al*. (2015), and NCBI (Sayers *et al*. 2021) databases. The conservation scores (or ranked phylogenetic profiling) were calculated using the protein sequence similarity between each gene in the human genome and its homologs in each of the species was measured by the bit score computed with BLASTP (Altschul *et al*. 1990). For each human gene and each species, we only considered the matched gene with the highest bit score. While other good approaches are available like reciprocal blast, our method has been widely used and worked well using human and other reference genomes (Bloch *et al*., 2020; Omar *et al*., 2018; Tsaban *et al*., 2021). To reduce the influence of random matches, the bit scores were set to 0 for matches with E-value > 1e-5. Bit score is known to be affected by the length of the reference protein. To eliminate the protein length effect, we normalize to protein length by dividing each bit score by the score of the reference protein against itself, resulting in values between 0 and 1 (Tabach *et al*. Nature 2013; Tabach *et al*. MSB 2013). Finally, the conservation scores were obtained by rank-normalizing the protein length-normalized bit scores across genes within each species, to control for the evolutionary distance between human and each species. These rank-normalized values range from 0 to 1, with higher values corresponding to higher levels of conservation. To examine whether the use of human (considered a relatively cancer-resistant species) as the reference affects the results, in a similar manner, we also repeated the above computation using a cancer prone species like house mouse as reference.

### Cancer resistance estimates

Since the cancer incidence in non-human species is unknown, we used two indirect methods to estimate the level of cancer resistance in a species. Let AW stand for adult weight and ML for maximum longevity of a species, we define the two cancer-resistance estimates/measures as follows:

*MLTAW measure:* log(ML^6^ × AW)
*MLCAW or “maximum longevity controlled for adult weight” measure:* Residue obtained by regressing out log(AW) from log(ML), using linear regression.
MLTAW and MLCAW were computed for 193 out of the 240 species for which both ML and AW data is available in the AnAge database (Tacutu *et al*. 2018). These 193 species are from various classes or taxonomy groups: 108 Mammalia (mammals), 55 Aves (birds), 18 Teleostei (ray-finned fishes), 7 Reptilia (reptiles), 1 Amphibia (amphibians), 1 Cephalaspidomorphi (jawless fishes), 1 Chondrichthyes (cartilaginous fishes), 1 Coelacanthi (lobe-finned fishes), 1 Holostei (bony fishes).

### Identification of cancer resistance-associated genes

To identify cancer resistance-associated genes (PC or NC genes), we computed the Pearson correlation coefficient between the conservation scores of each gene and each of the two cancer-resistance estimates (MLTAW and MLCAW) after proper transformation (described above). Pearson correlation was chosen (instead of Spearman’s correlation) so as to reduce the number of ties in further GSEA analysis for pathway enrichment. The robust identification of PC/NC genes is independent of the correlation measure used (see Supp. Note for details). Among the genes with Benjamini-Hochberg-adjusted P values (FDR) less than 0.1 or 0.01, those with correlation estimates > 0 are defined as PC genes while those with correlation estimates < 0 are NC genes. This analysis was done for all species or within certain groups of species. PC and NC genes were identified separately based on each of the two cancer-resistance estimates (MLTAW/MLCAW).

### Cancer resistance predictor

Since higher conservation scores of the PC genes corresponds to higher level of cancer resistance, and vice versa for the NC genes, we define a cancer resistance (CR) score for each species as follows:

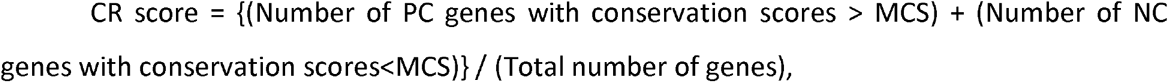

where MCS is the median conservation score of all genes in a species. PC and NC genes are chosen for FDR<0.1. We also repeat this analysis for different thresholds (some other quantile other than median) or FDR thresholds for robustness studies. Total number of genes = 20,076 in our analysis when we used human as reference.

Cross-validation analysis was mainly done in a leave-one-out manner. For each test sample, we identify PC and NC genes on the training set and predict CR scores on the test set.

For robustness tests, we also did a two-fold cross validation, i.e. identify PC and NC genes on the training group and test the accuracy of the CR predictions on the left-out group. We also do cross-validation by leaving out an entire group of species and identify PC and NC genes from the remaining species, and testing on the left-out group. For the all-species analysis we left one class out (for different classes) and for the mammalian analysis we left one order out (for different orders).

### Modifications of cancer resistance predictions

We explored the prediction of cancer resistance using only PC genes or NC genes as follows: CR score = (Number of PC genes with conservation scores > MCS) / (Total number of genes), or CR score = (Number of NC genes with conservation scores < MCS) / (Total number of genes). We also predicted cancer resistance using either human tumor suppressor genes (TSGs) and oncogenes obtained from the Cancer Gene Census dataset from the COSMIC database (Forbes *et al*. 2015). Specifically, we used either TSGs alone, or oncogenes alone, or TSGs combined with oncogenes to compute the CR score: (Number of TSGs, or oncogenes, or combined > MCS) / (Total number of genes), where MCS is the median conservation score of all genes in a species.

### Pathway, cancer driver gene and other cancer-related gene set enrichment analysis

The biological pathway annotation data was downloaded from the Reactome database (Jassal *et al*. 2020). The sets of curated human oncogenes and TSGs were obtained from the Cancer Gene Census dataset from the COSMIC database (Forbes *et al*. 2015). Significant markers reported in various GWAS studies linked to human cancers were collected from the EBI GWAS Catalog database (Buniello *et al*. 2019) using the keyword “cancer” as the phenotypes/traits. Variants in stronger linkage disequilibrium (LD) (with D’ ≥ 0.8 and r2 ≥ 0.3) with the GWAS associated markers (within 500K base pairs in each side) in loci, replicated in more than one studies, were selected using the R package, LDlinkR (Myers *et al*. 2020), and genes containing such variants were selected. The sets of genes whose knockout can result in various cancer-related phenotypes in mice were obtained from the MGI database (Bult *et al*. 2019). Specifically, we selected the genes for which the allele attributes are “Null/knockout”, and the corresponding phenotype terms are among “increased cancer incidence”, “decreased cancer incidence”, “increased cancer latency”, and “decreased cancer latency”. The set of loss-of-function genes identified in canine transmissible venereal tumors were obtained from (Murchison *et al*. 2014). The enrichment for each of the biological pathways and cancer-related gene sets based on the MLTAW or MLCAW correlation results was tested with gene set enrichment analysis (GSEA, Subramanian *et al*. 2005).

### LOEUF score analysis for PC and NC genes

The LOEUF scores of all human genes were obtained from (Karczewski *et al*. 2020). Lower LOEUF scores correspond to less tolerance to loss-of-function genomic variations in humans. The LOEUF scores of the PC (or NC) genes we identified were compared with each other or to the rest of the genes in the human genome with Wilcoxon rank-sum tests. Given that a homogenous adjusted P value cutoff produced drastically different numbers of significant genes from different analysis (e.g. with adjusted P<0.05, there are more than 2800 significant genes from the primate MLTAW correlation analysis, but only 78 from the rodent MLTAW correlation), here, the PC and NC genes were selected instead based on the criterion of “expected number of false discoveries (Gordon *et al*. 2007) smaller than 1”. But if this criterion results in fewer than 100 genes, then a less stringent criterion of “expected number of false discoveries smaller than 5” was used. These sets of PC and NC genes are given in Table S6A.

### Analysis of the genes associated with lifetime cancer risk across human tissues

The data on the lifetime cancer risk in each of the human tissue/organ sites were obtained from the SEER database (SEER program, 2018), and RNA-seq data of normal human tissue samples across multiple tissue types were downloaded from the GTEx database (Lonsdale *et al*. 2013). For each gene, we computed its median expression level in each tissue type, then computed the Spearman’s correlation coefficient between the median expression value and the lifetime cancer risk across tissue types. All the genes were ranked by the Spearman’s correlation coefficient, and the enrichment of the PC or NC genes for genes associated with lifetime cancer risk across human tissues was tested with GSEA. The criterion for identifying PC and NC genes is the same as that described in the section above (“LOEUF score analysis for PC and NC genes”).

### Selection of subsets of PC/NC genes with high relevance to human cancers based on multiple criteria

For each of the PC/NC genes from the various analyses (at FDR<0.1), we look for supporting evidence from many of the different analyses described in the manuscript. Evidence considered are if a gene is: (a) a PC or NC gene (at FDR<0.1) for the all-species, mammals-only, birds-only analysis using both the estimates; (b) human oncogene or tumor suppressor; (c) whose knockout causes early cancer incidence or early cancer onset in mice; (d) is a loss-of-function gene in CTVT; (e) GWAS gene associated with human cancers; (f) expressed mutated genes in single-cell phylogeny of a mouse melanoma model. We then rank each gene by the number of times of support in Table S8.

## CODE/DATA AVAILABILITY

The code used for cancer resistance prediction and pathway analysis is made available in (https://hpc.nih.gov/~nairnu/species_cancer_resistance.zip) for the sake of reproducibility.

## ACKNOWLEDGEMENTS

This research was supported in part by the Intramural Research Program of the National Institutes of Health (NIH), National Cancer Institute and the Center for Cancer Research. This work utilized the computational resources of the NIH HPC Biowulf cluster (http://hpc.nih.gov). We thank Dr. Xin Wei Wang for his helpful comments on this work.

## CONFLICT OF INTEREST STATEMENTS

The authors declare no conflict of interest.

## Notes

### Competing Interest Statement

The authors have declared no competing interest.

