## Supplementary notes and supplementary figures for "Cross-species identification of cancer-resistance associated genes uncovers their relevance to human cancer risk"

### Supplementary notes and supplementary figures for the paper titled "Cross-species identification of cancer-resistance associated genes uncovers their relevance to human cancer risk"

Nishanth Ulhas Nair[1,*,#], Kuoyuan Cheng[1,2,*,#], Lamis Naddaf[3,*], Elad Sharon[3], Lipika R. Pal[1], Padma S. Rajagopal[4], Irene Unterman[3], Kenneth Aldape[5], Sridhar Hannenhalli[1], Chi-Ping Day[6], Yuval Tabach[3,#], Eytan Ruppin[1,#]

1. Cancer Data Science Laboratory (CDSL), National Cancer Institute (NCI), National Institutes of Health (NIH), Bethesda, MD, USA.
2. Center for Bioinformatics and Computational Biology, University of Maryland, College Park, MD, USA.
3. Department of Developmental Biology and Cancer Research, Institute of Medical Research - Israel-Canada, The Hebrew University of Jerusalem, Jerusalem 9112102, Israel.
4. Section of Hematology/Oncology, Department of Medicine, The University of Chicago, Chicago, IL, USA.
5. Laboratory of Pathology, National Cancer Institute (NCI), National Institutes of Health (NIH), Bethesda, MD, USA.
6. Laboratory of Cancer Biology and Genetics, National Cancer Institute (NCI), National Institutes of Health (NIH), Bethesda, MD, USA.

* These authors contributed equally to this work as co-first authors.

#### SUPPLEMENTARY NOTES

##### Short review of cancer resistance mechanisms in different species

Different species have independently evolved unique cancer resistance mechanisms. Repression of somatic telomerase activity, and replicative senescence are important tumor-suppressing mechanisms evolved in species greater than approximately 10 kilograms (Gorbunova *et al.* 2014). Cells of smaller but relatively long-lived animals are reported to have slower proliferation in culture (Tian *et al.* 2018). There have been a few reports of more efficient DNA repair in cancer-resistant and long-lived animals (Tollis *et al.* 2017; Seluanov *et al.* 2018). African elephants, the largest land mammals, have 19 extra retrogene copies of the tumor suppressor gene *TP53* and are more sensitive to *TP53*-mediated DNA damage response (Abegglen *et al.* 2015; Sulak *et al.* 2016). The remarkable cancer resistance of naked mole rat has been previously partly attributed to the production of high molecular mass hyaluronan (HMM-HA) (Buffenstein, 2008; Tian *et al.* 2013) but this mechanism has been questioned in a recent study (Hadi *et al.* 2020). Blind mole rats also have abundant HMM-HA and increased interferon-β expression that contribute to cancer resistance (Gorbunova *et al.* 2012; Tian *et al.* 2013; Seluanov *et al.* 2018). Various large-bodied whales do not have additional *TP53* copies, and their cancer resistance mechanisms are not clearly understood (Seluanov *et al.* 2018; Keane *et al.* 2015).

##### Cancer resistance prediction in birds and teleost fishes

Using leave-one-out cross-validation (LOOCV) we find a significant positive correlation between the predicted cancer resistance (CR) scores and MLTAW all the bird species (Spearman's $\rho = 0.43$, $P = 0.00094$, Fig. S3A) though this is weaker than the corresponding predictions obtained by learning on all species (Spearman's $\rho = 0.57$, $P = 6.4e-6$, Fig. S15A, using LOOCV). The correlation is stronger for the order Passeriformes (Spearman's $\rho = 0.79$, $P = 0.0012$, Fig. S3B) for which we have the largest number of samples. Among Passeriformes, the highest CR scores are obtained for American crow (Fig. S3B). The MLCAW measure did not yield any PC/NC genes ($FDR < 0.1$) for birds, and hence no CR predictor could be built. We could not identify any PC/NC genes at FDR <

0.1 for teleost fishes (for both MLTAW and MLCAW measures) probably because of a small sample size (n=18); and hence we could not however build a cancer resistance predictor for them.

A detailed pathway enriched analysis for mammals, birds, and teleost fishes are provided in Figs. 3A, S12C. Many pathways show group-specific enrichment. For example, many cell cycle and DNA repair-related pathways are enriched by the PC genes in mammals, but not or to a much lesser extent in birds or teleost fishes; complement activation is enriched by PC genes in teleost fishes, but by NC genes in mammals or birds (Figs. 3A, S12C, Table S3). Bird PC genes are uniquely enriched for certain processes including fatty acid and amino acid metabolism and PI3K-AKT signaling pathway despite sharing interleukin, interferon signaling and mRNA transcription with mammals (Figs. 3A, S12C). GPCR signaling is commonly enriched by the NC genes based on MLCAW in all three groups (Fig. 3A, S12C, Table S3).

**Control and robustness analysis**

Random control experiments

Random controls experiments for predicting cancer resistance were done for using all species. We chose random PC/NC genes with the same size as the actual PC/NC genes identified from the all-species analysis at FDR < 0.1. We can predict cancer resistance using these genes. We do this for 1000 iterations and the empirical P-value is computed. We see that they are not correlated in comparison to the 'true' correlation obtained using the actual PC/NC genes (randomization test  P < 0.001). The results for MLTAW/MLCAW are shown in Fig. S4.

PC/NC genes and cancer resistance predictors are robust to the method of correlation used.

To identify cancer resistance-associated genes (PC/NC genes) for all species, we used Pearson correlation between the conservation scores of each gene and the cancer-resistance estimates (MLTAW and MLCAW) across all species. Pearson correlation coefficient was used (instead of Spearman) in order to reduce the number of ties which will affect the gene set enrichment analysis (GSEA). We now show that the robust identification of PC/NC genes is possible even if we use Spearman's correlation instead of Pearson. To do this, we recomputed PC/NC genes using Spearman's correlation and compared it to those obtained using Pearson's correlation (FDR <

0.1). Using Fisher's exact test, we get a significant overlap in comparing cancer resistance-associated genes obtained using Pearman and Spearman's correlation (PC genes: Odds-ratio/OR = 111.64, P < 2.2e-16 for MLTAW and OR = 94.22, P < 2.2e-16 for MLCAW; NC genes: OR = 126.52, P < 2.2e-16 for MLTAW and OR = 184.78, P < 2.2e-16 for MLCAW). (Note: whenever the enrichment test software shows P = 0, we write as P < 2.2e-16.) Cancer-resistance (CR) predictors computed from PC/NC genes identified using Pearson or Spearman's correlation works very similar across all species (Pearson-based PC/NC identification: MLTAW ρ=0.44, P=1.32e-10, MLCAW ρ=0.51, P=2.31e-14; Spearman's-based PC/NC identification: MLTAW ρ=0.43, P=3.65e-10, MLCAW ρ=0.51, P=5.16e-14; using LOOCV or 'leave-one-out cross-validation').

PC/NC genes and cancer resistance predictors are robust to the choice of reference species

We used the human genome as a reference, for computing gene conservation matrix. To check if our analysis is robust to changes in reference, we recomputed the gene conservation matrix using the house mouse (Mus Musculus) genome or thirteen-lined ground squirrel (Ictidomys tridecemlineatus) genome as a reference. House mouse was chosen as it is considered a cancer-prone species in comparison to humans (which are relatively cancer resistant). In the all-species analysis, the PC/NC genes obtained using mouse or squirrel genome as reference are extremely similar to the PC/NC genes obtained using humans as reference. Using Fisher's exact test, we get a significant overlap in comparing the PC genes obtained using humans and mouse/squirrel genomes as reference (mouse: OR = 14.26, P < 2.2e-16 for MLTAW and OR = 30.69, P < 2.2e-16 for MLCAW; squirrel: OR = 33.78, P < 2.2e-16 for MLTAW and OR = 55.31, P < 2.2e-16 for MLCAW). Similarly, we get a significant overlap in comparing the NC genes obtained using humans and mouse/squirrel genomes as reference (mouse: OR = 11.01, P < 2.2e-16 for MLTAW and OR = 27.76, P < 2.2e-16 for MLCAW; squirrel: OR = 24.29, P < 2.2e-16 for MLTAW; OR = 57.68, P < 2.2e-16 for MLCAW). (Note: whenever the enrichment test software shows P = 0, we write as P < 2.2e-16.) Similarly, cancer-resistance (CR) predictors computed from a gene conservation matrix which uses mouse or squirrel genomes as reference, show good prediction results between CR scores and cancer-resistance estimates; similar to what was obtained using human as reference (Fig. S5; LOOCV). We also see that humans are predicted to be relatively cancer resistant as expected (Fig. S5).

Using two-fold cross validation instead of LOOCV

We also did a two-fold cross validation (instead of LOOCV) for predicting cancer-resistance scores, i.e. identify PC and NC genes on the training group and test the accuracy of the CR predictions on the left-out group. We see that our results using two-fold cross-validation is quite similar to that obtained by LOOCV in the all-species analysis (Fig. S6 in comparison to Figs. 2A, S2).

Our results are robust to changes in FDR criteria and thresholds used

We predicted cancer-resistance scores (CR) by altering various parameters. Our original predictor as described in the manuscript uses PC genes and NC genes which are significantly associated with cancer resistance at FDR < 0.1. We now show that our CR predictor is robust to changes in FDR thresholds from 0.1 to 0.01 or 0.2 (Figs. S7A,B). The original predictor also computes the number of PC genes whose conservation score > median conservation score; and the number of NC genes whose conservation score < median conservation score. We also show that the CR predictor is robust to altering the thresholds from median conservation score to top and bottom 33 percentile of the conservation scores for PC and NC genes respectively (Fig. S7C). The same analysis was also done using top and bottom 20 percentile (Fig. S7D).

Alternative predictors using either PC or NC genes

For the original predictor which uses both PC and NC genes, our cancer resistance (CR) score was measured using the following equation:

*Original predictor:* CR score = {(No. of PC genes > MCS) + (No. of NC genes < MCS)} / (Total no. of genes)

where MCS is the median conservation score of all genes in a species; PC and NC genes are chosen for FDR < 0.1.

Now to test the individual contribution of using PC-only and NC-only genes to predict a good cancer-resistance estimate, we build alternative predictors as follows:

*PC-only predictor:* CR score = (No. of PC genes > MCS) / (Total no. of genes)

*NC-only predictor:* CR score = (No. of NC genes < MCS) / (Total no. of genes)

We then compare the PC-only and NC-only predictor with the original predictor in Fig. S8 for the all-species, Mammalia (mammals), and Aves (birds) analysis for both MLTAW and MLCAW measures. We see that both PC and NC genes have significant individual and comparable contributions to predict cancer resistance.

**Cancer resistance prediction within specific mammalian orders**

Using the predicted CR scores learnt from all mammalian species (in LOOCV), we further tested its association with cancer-resistance estimates for various orders with the class mammalia: rodentia, primates, carnivora, artiodactyla, cetacea, chiroptera. We are able to predict at least one of the two cancer-resistance estimates effectively in rodentia, primates, carnivora, and chiroptera; but not for artiodactyla, cetacea (Figs. S9A,B). Just looking at rodents, we are able to get good CR scores in known cancer resistant species like the naked mole rat and low CR scores for cancer prone species like the house mouse (Fig. S9C). Among primates, we see that animals like chimpanzees and gorillas are predicted to be cancer resistant (Fig. S9D). Among carnivores, we see that Steller sea lion and California sea lion have high CR scores (Fig. S9E). Among bats, we again see that species like the Brandt's bat which are known to live long for their body size are predicted to have high CR scores (Fig. 2D). Little brown bats are also seen to have high CR scores (Fig. 2D).

**Adaptation to different oxygen levels and cancer resistance**

In our study, we have not explicitly considered factors potentially linked to variations in cancer resistance that are not reflected through body size and lifespan. These factors are more difficult to quantify with limited data available. One example is the adaptation to different oxygen concentrations and oxidative stress levels. Reactive oxygen species (ROS) levels have a complicated role in cancers, although one of the effects of ROS is DNA damage, which is linked to cancer development (Waris *et al.* 2006), and tolerance to hypoxia is also associated with cancer resistance, as evident in several well-known cancer resistant species including the naked mole rat and certain bats (Ilacqua *et al.* 2017; Hammarlund *et al.* 2018). This was not considered in our analysis due to the challenge to quantitate hypoxia resistance for each species, which may underlie some notable outliers in our cancer resistance predictions. For example, the predicted gene conservation-based CR score was high for the small and short-lived star nosed mole (55 grams, 2.5 years; Figs. 2A, S2), which largely lives underground and is hypoxia-tolerant (McIntyre *et al.* 2002). When more phenotypic data across species become available in the future, further studies are required to refine and update our findings here.

#### SUPPLEMENTARY FIGURES

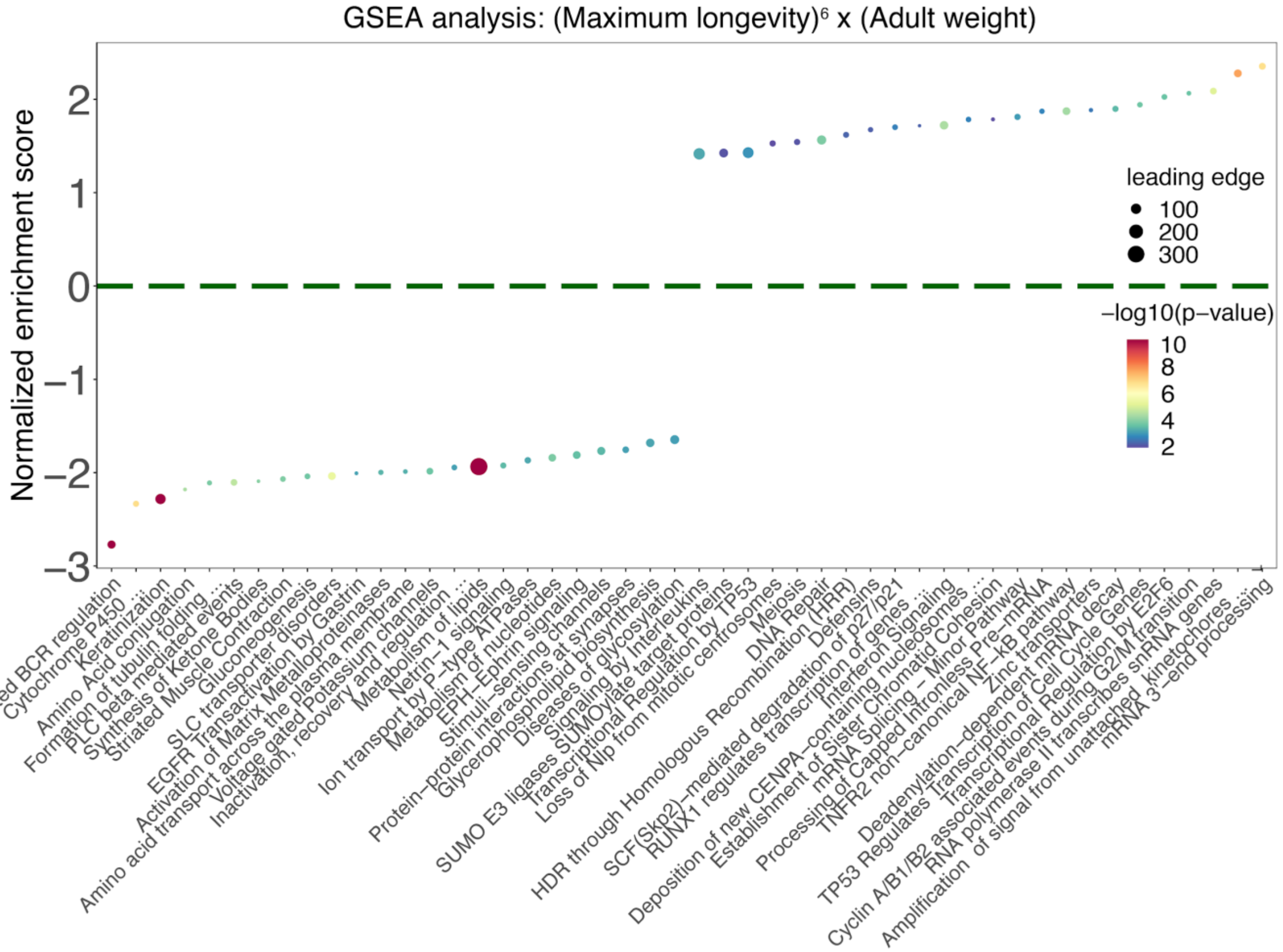


***Figure S1:*** *Summary of the top significantly enriched pathways (adjusted P<0.1) by the genes whose conservation scores are correlated with cancer-resistance estimates, using gene set enrichment analysis (GSEA) with gene set annotations from the Reactome database. The cancer-resistance estimates used is MLTAW or '(Maximum longevity)$^6$ x (adult weight)'. Normalized enrichment score is plotted on the Y-axis, where positive values correspond to enrichment by the positively correlated (PC) genes and negative values correspond to enrichment by the negatively correlated (NC) genes. The dot color represents the significance of the enrichment (negative log10 GSEA P value), and the dot size represents the number of genes in the "leading edge", i.e. the set of genes that are enriched in a pathway. For the sake of clarity, only a subset of the enriched*

*pathways (FDR<0.1) are shown and long pathway names have been shortened (using "..."). The complete pathway enrichment results are given in Table S3A.*

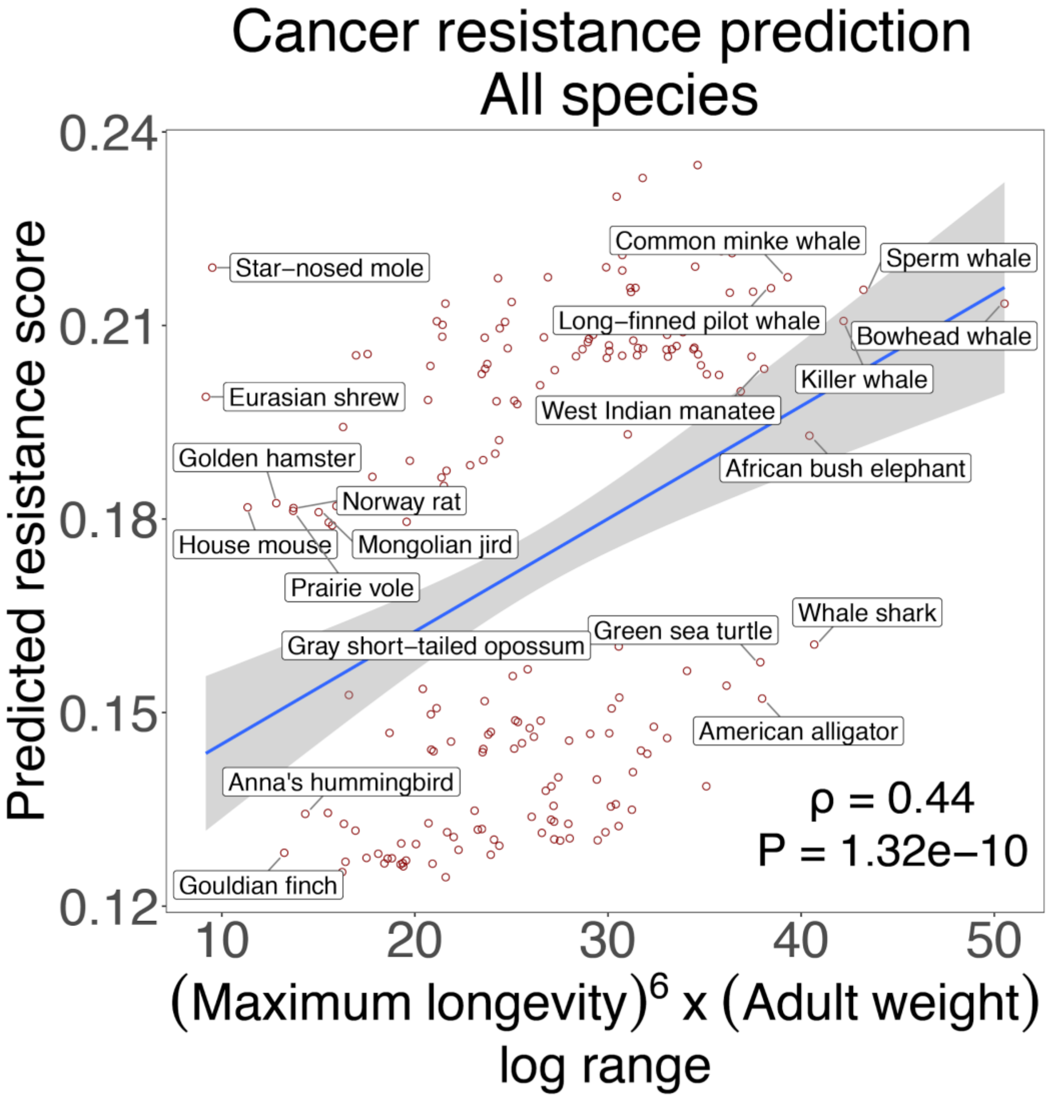


***Figure S2:*** *Scatter plots showing the correlation between the predicted cancer resistance (CR) scores computed based on gene conservation and for the cancer-resistance estimate MLTAW or '(Maximum longevity)$^6$ x (adult weight)', with leave-one-out cross-validation. Results for all species. Species with the top and bottom 10% MLTAW values are labeled by their common names for the sake of clarity. Spearman's ρ and p-values (P) are shown.*

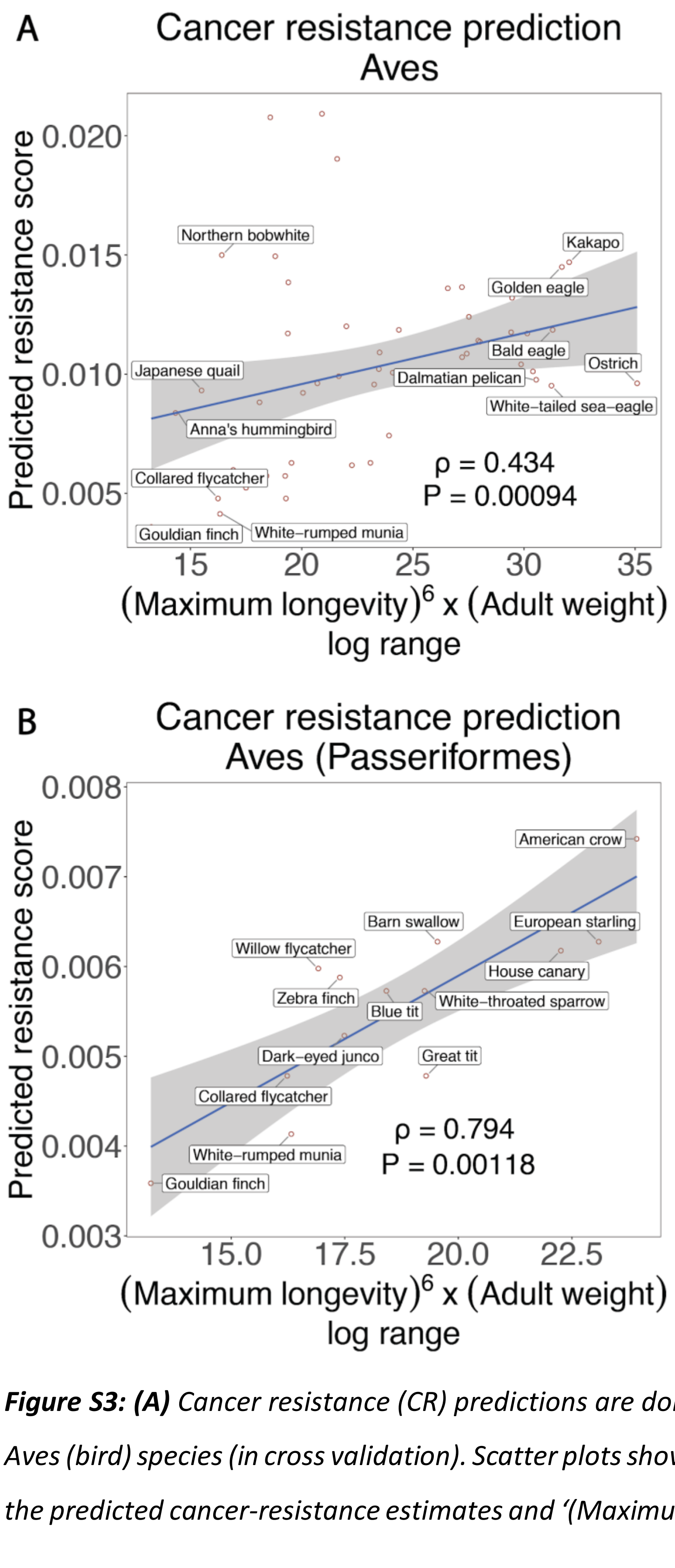


***Figure S3: (A)*** *Cancer resistance (CR) predictions are done by identifying PC/NC genes using only Aves (bird) species (in cross validation). Scatter plots showing the Spearman's correlation between the predicted cancer-resistance estimates and '(Maximum longevity)$^6$ x (adult weight)' or MLTAW*

*is shown. Only species names for the top and bottom 10 percentile of the MLTAW measure are labelled for display clarity.* ***(B)*** *Scatter plots using the predicted scores in (A) are shown for only order Passeriformes (n=14) within the bird species. Spearman's ρ and p-values (P) are reported for (E,F). No CR predictor was built for birds using the MLCAW measure as we did not identify any PC/NC genes at (FDR < 0.1) for this measure.*

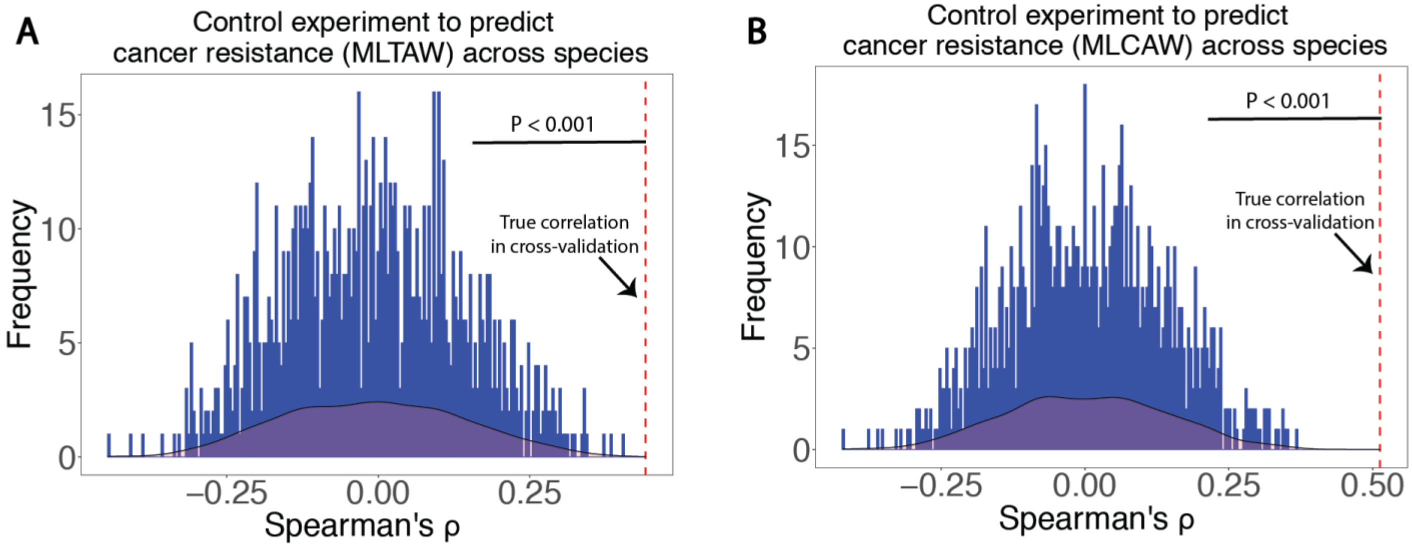


***Figure S4:*** *Random controls experiments for predicting cancer resistance using all species. We chose random PC/NC genes with the same size as the actual PC/NC genes identified from the all species analysis at FDR < 0.1. We can predict cancer resistance using these genes. We do this for 1000 iterations and the empirical P-value is computed. We see that they are not correlated in comparison to the 'true' correlation obtained using the actual PC/NC genes (randomization test P < 0.001). Cancer-resistance estimates used are: (A) MLTAW; (B) MLCAW.*

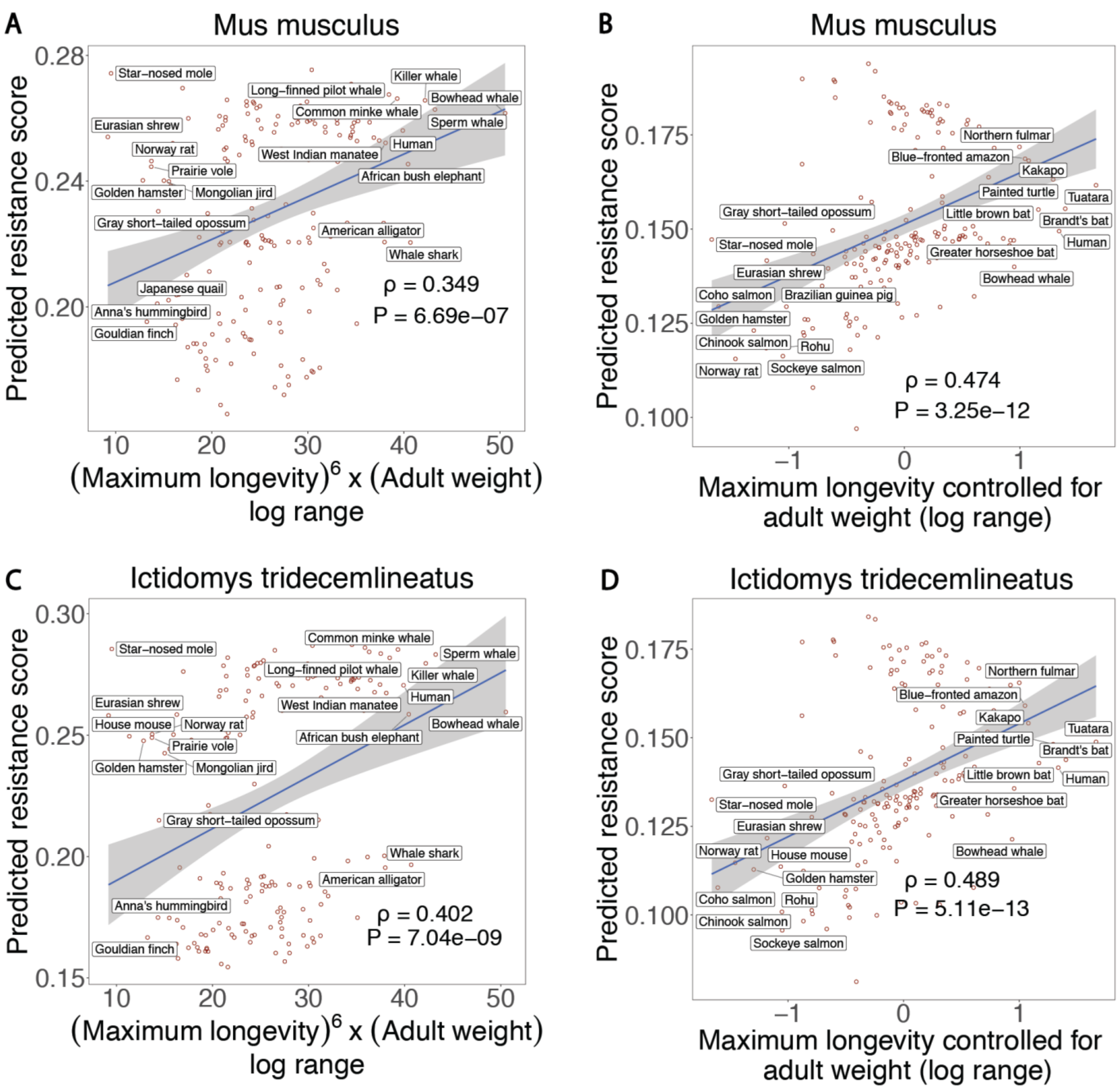


***Figure S5:*** *Instead of human reference to compute gene conservation scores, we use Mus musculus (house mouse) and thirteen-lined ground squirrel (Ictidomys tridecemlineatus) as reference. We predict cancer resistance in this all-species analysis. Scatter plots along with the Spearman's correlation between the predicted cancer-resistance estimates (in cross-validation, LOOCV) and the cancer-resistance estimates like* ***(A)*** *MLTAW or '(Maximum longevity)$^6$ x (adult weight)' for mouse;* ***(B)*** *MLCAW or 'Maximum longevity controlled for adult weight' for mouse;*

***(C)*** *MLTAW for squirrel;* ***(D)*** *MLCAW for squirrel, are shown. Both Spearman's ρ and p-values (P) are reported. Only species names for the top and bottom 5 percentile of the MLTAW/MLCAW measures are labelled for display clarity. We see that humans are predicted to be relatively cancer resistant as expected. The results obtained are quite similar to the corresponding results using human reference.*

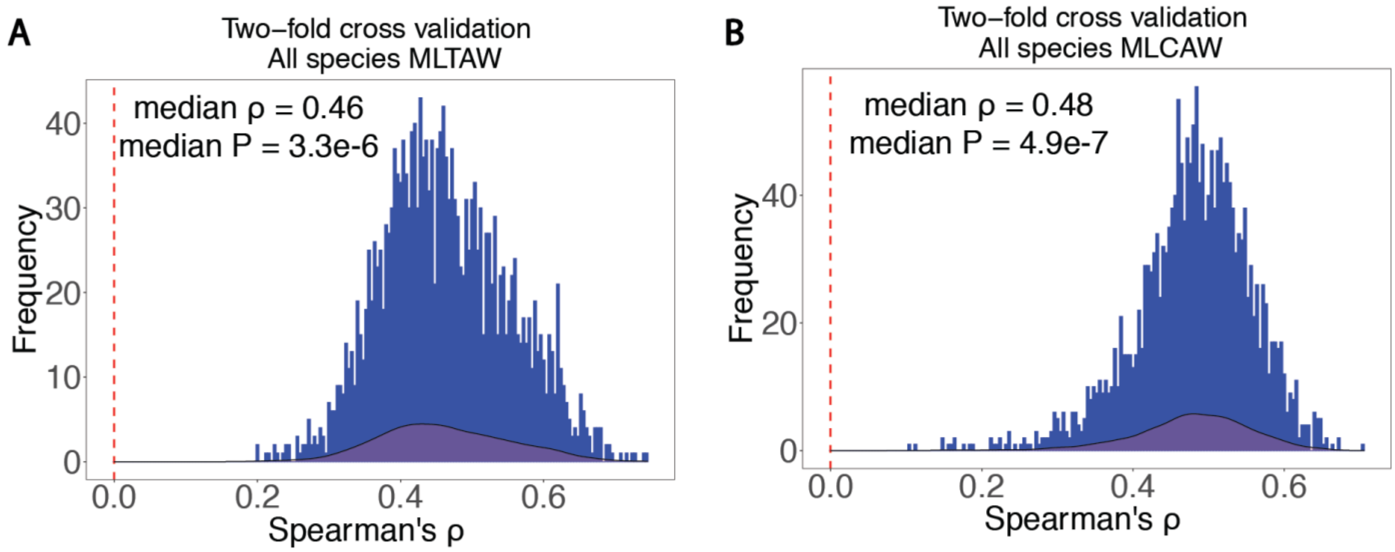


***Figure S6:*** *Plots show the distribution/frequency of Spearman's ρ between the predicted cancer resistance (CR) scores computed based on gene conservation and each of the two cancer-resistance estimates: (A) MLTAW, i.e. (Maximum longevity)$^{6}$ x (adult weight); (B) MLCAW i.e. maximum longevity controlled for adult weight), using two-fold cross-validation (instead of LOOCV). The two-fold cross validation was carried out 1000 times (2000 data points). Median Spearman's ρ and p-values (P) are mentioned.*

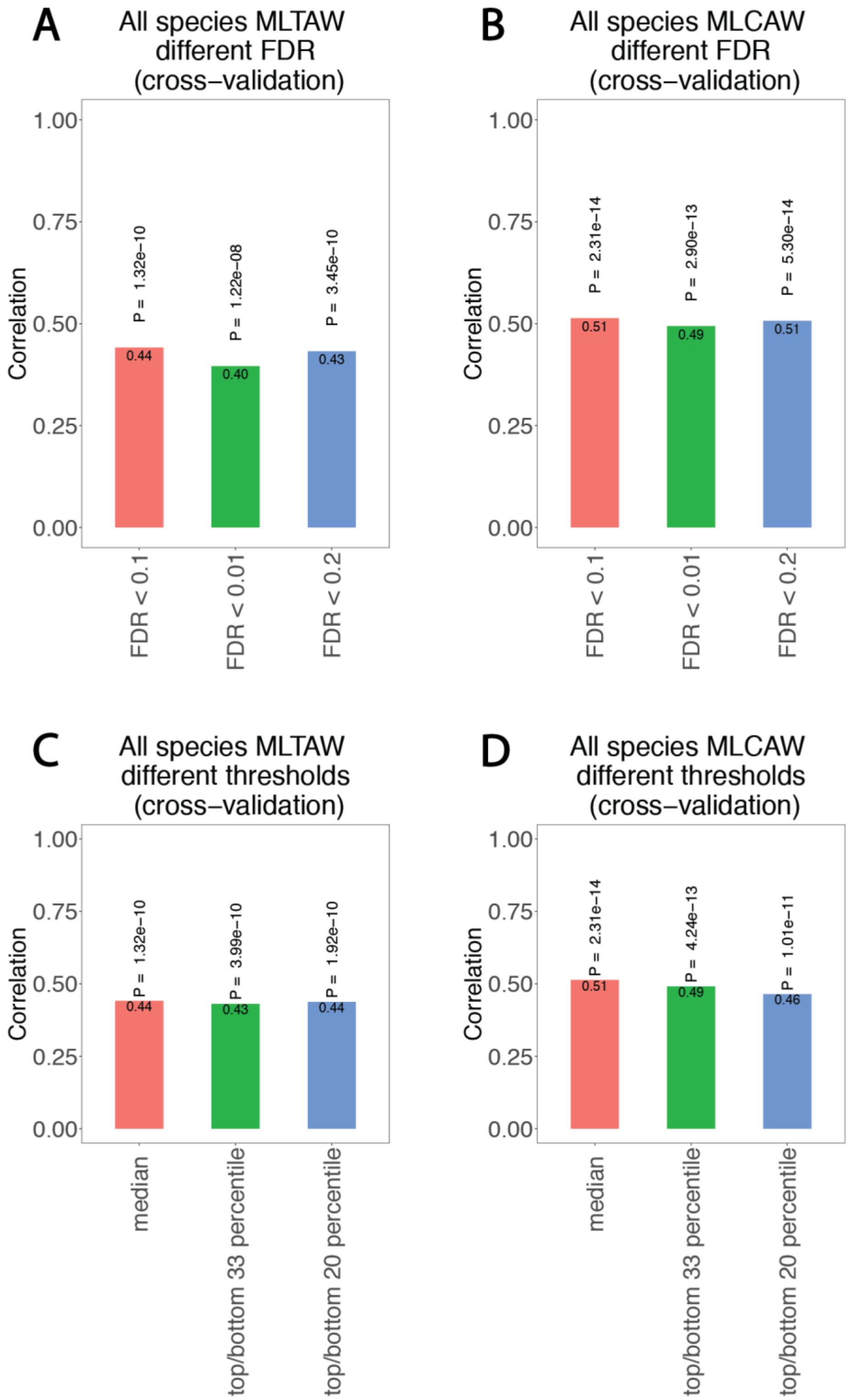
A
All species MLTAW different FDR (cross−validation)
Correlation
1.00
0.75
0.50
0.25
0.00
P = 1.32e−10
P = 1.22e−08
P = 3.45e−10
0.44
0.40
0.43
FDR < 0.1
FDR < 0.01
FDR < 0.2
B
All species MLCAW different FDR (cross−validation)
P = 2.31e−14
P = 2.90e−13
P = 5.30e−14
0.51
0.49
0.51
C
All species MLTAW different thresholds (cross−validation)
P = 1.32e−10
P = 3.99e−10
P = 1.92e−10
0.44
0.43
0.44
median
top/bottom 33 percentile
top/bottom 20 percentile
D
All species MLCAW different thresholds (cross−validation)
P = 2.31e−14
P = 4.24e−13
P = 1.01e−11
0.51
0.49
0.46

***Figure S7:*** *We predicted cancer-resistance (CR) scores by altering various parameters. CR predictors for different FDR thresholds (0.01, 0.01, or 0.2) are shown using* ***(A)*** *MLTAW; and* ***(B)*** *MLCAW measures. The original predictor also computes the number of PC genes whose conservation score > median conservation score; and the number of NC genes whose conservation score < median conservation score. CR predictor results are shown to be robust by altering the thresholds from median conservation score to top and bottom 33/20 percentile of the conservation scores for PC and NC genes respectively (****C,D****). Both Spearman's ρ and p-values (P) are reported.*

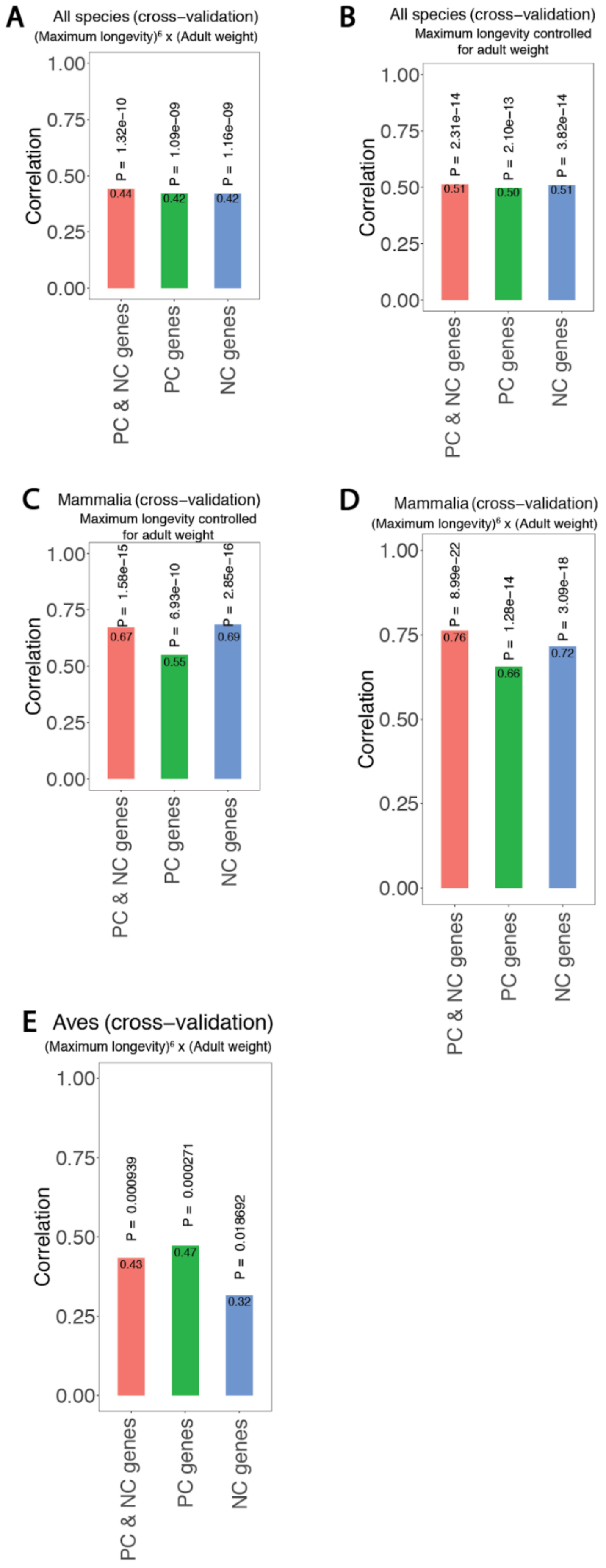
A
All species (cross−validation)
(Maximum longevity)⁶ x (Adult weight)
Correlation
P = 1.32e−10
P = 1.09e−09
P = 1.16e−09
0.44
0.42
0.42
PC & NC genes
PC genes
NC genes
B
All species (cross−validation)
Maximum longevity controlled for adult weight
P = 2.31e−14
P = 2.10e−13
P = 3.82e−14
0.51
0.50
0.51
C
Mammalia (cross−validation)
Maximum longevity controlled for adult weight
P = 1.58e−15
P = 6.93e−10
P = 2.85e−16
0.67
0.55
0.69
D
Mammalia (cross−validation)
(Maximum longevity)⁶ x (Adult weight)
P = 8.99e−22
P = 1.28e−14
P = 3.09e−18
0.76
0.66
0.72
E
Aves (cross−validation)
(Maximum longevity)⁶ x (Adult weight)
P = 0.000939
P = 0.000271
P = 0.018692
0.43
0.47
0.32

***Figure S8:*** *We predicted cancer-resistance (CR) scores by using both PC and NC genes (PC & NC genes); PC genes only; NC genes only. Results for MLTAW or '(Maximum longevity)$^{6}$ x (adult weight)'; and MLCAW or 'Maximum longevity controlled for adult weight' are shown for the all-species analysis* ***(A,B)****, Mammalia-only analysis* ***(C,D)****, and Aves-only* ***(E)*** *analysis. MLCAW analysis is not shown for birds as we could not identify PC/NC genes at FDR < 0.1; and therefore could not build a CR predictor.*

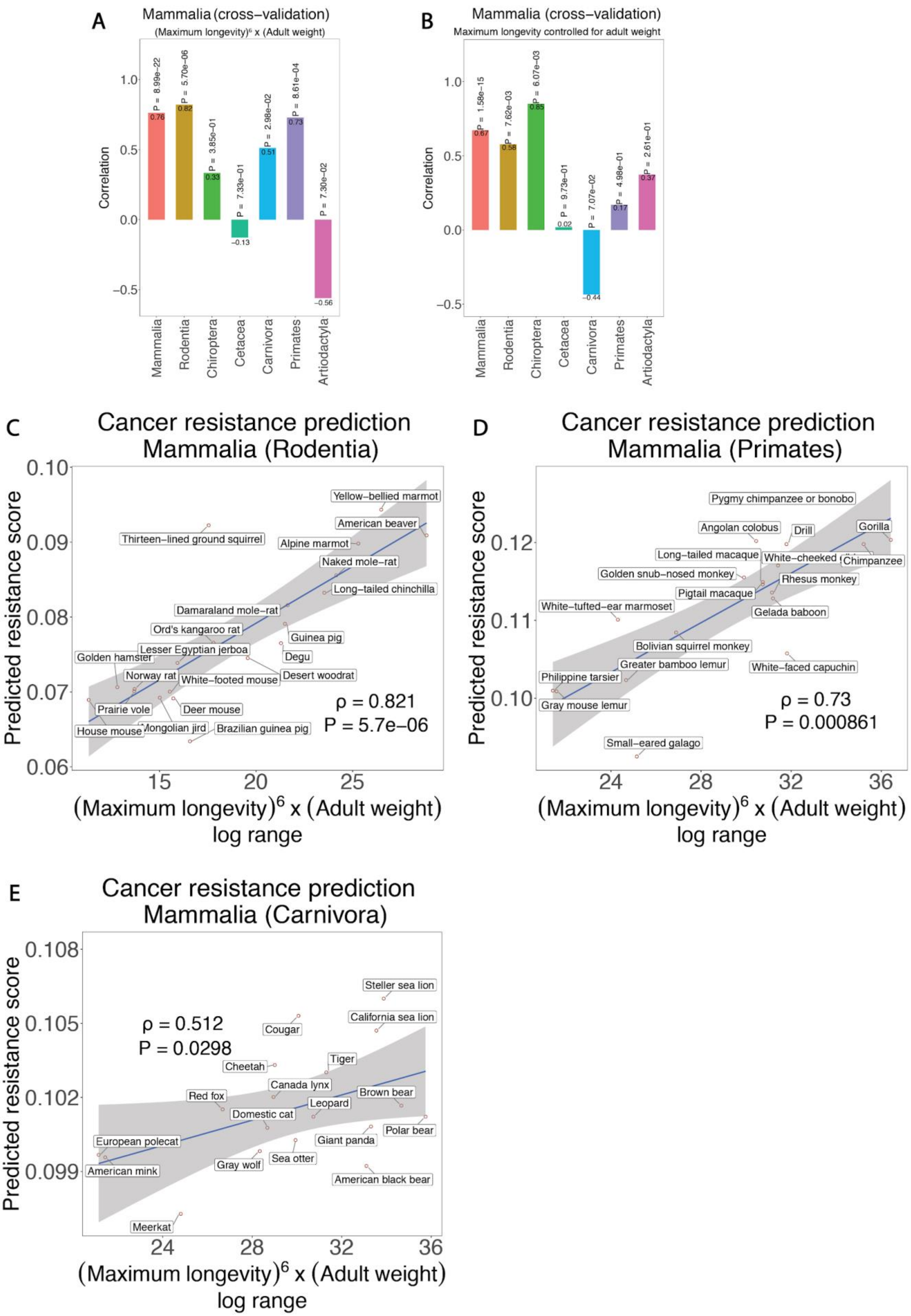
A
Mammalia (cross−validation)
(Maximum longevity)⁶ x (Adult weight)
Correlation
P = 8.99e−22
P = 5.70e−06
P = 3.85e−01
P = 7.33e−01
P = 2.98e−02
P = 8.61e−04
P = 7.30e−02
0.76
0.82
0.33
−0.13
0.51
0.73
−0.56
Mammalia
Rodentia
Chiroptera
Cetacea
Carnivora
Primates
Artiodactyla
B
Mammalia (cross−validation)
Maximum longevity controlled for adult weight
Correlation
P = 1.58e−15
P = 7.62e−03
P = 6.07e−03
P = 9.73e−01
P = 7.07e−02
P = 4.98e−01
P = 2.61e−01
0.67
0.58
0.85
0.02
−0.44
0.17
0.37
Mammalia
Rodentia
Chiroptera
Cetacea
Carnivora
Primates
Artiodactyla
C
Cancer resistance prediction
Mammalia (Rodentia)
Predicted resistance score
(Maximum longevity)⁶ x (Adult weight)
log range
Yellow−bellied marmot
American beaver
Thirteen−lined ground squirrel
Alpine marmot
Naked mole−rat
Long−tailed chinchilla
Damaraland mole−rat
Ord's kangaroo rat
Guinea pig
Lesser Egyptian jerboa
Golden hamster
Degu
Norway rat
White−footed mouse
Desert woodrat
Prairie vole
Deer mouse
House mouse
Mongolian jird
Brazilian guinea pig
ρ = 0.821
P = 5.7e−06
D
Cancer resistance prediction
Mammalia (Primates)
Predicted resistance score
(Maximum longevity)⁶ x (Adult weight)
log range
Pygmy chimpanzee or bonobo
Angolan colobus
Drill
Gorilla
Long−tailed macaque
White−cheeked
Chimpanzee
Golden snub−nosed monkey
Rhesus monkey
Pigtail macaque
White−tufted−ear marmoset
Gelada baboon
Bolivian squirrel monkey
Greater bamboo lemur
White−faced capuchin
Philippine tarsier
Gray mouse lemur
Small−eared galago
ρ = 0.73
P = 0.000861
E
Cancer resistance prediction
Mammalia (Carnivora)
Predicted resistance score
(Maximum longevity)⁶ x (Adult weight)
log range
Steller sea lion
California sea lion
Cougar
ρ = 0.512
P = 0.0298
Cheetah
Tiger
Canada lynx
Red fox
Brown bear
Leopard
Domestic cat
Polar bear
European polecat
Giant panda
Sea otter
Gray wolf
American mink
American black bear
Meerkat

***Figure S9:*** *Cancer resistance predictions were done on the entire mammalian species (LOOCV, by learning PC/NC genes from mammals). Using these predictions, Spearman's correlation (ρ and p-values) for different orders (sub-groups) of mammals: Rodentia (rodents), Chiroptera (bats), Cetacea (aquatic mammals like whales), Carnivora (carnivores), Primates, Artiodactyla (even-toed hoofed mammals) are shown for (A) MLTAW or '(Maximum longevity)$^6$ x (adult weight)' and (B) MLCAW or 'Maximum longevity controlled for adult weight'. Scatter plots showing the Spearman's correlation between the predicted cancer-resistance estimates and the MLTAW cancer-resistance estimate for some of the orders are shown in (C-E). Spearman's ρ and p-values (P) are reported.*

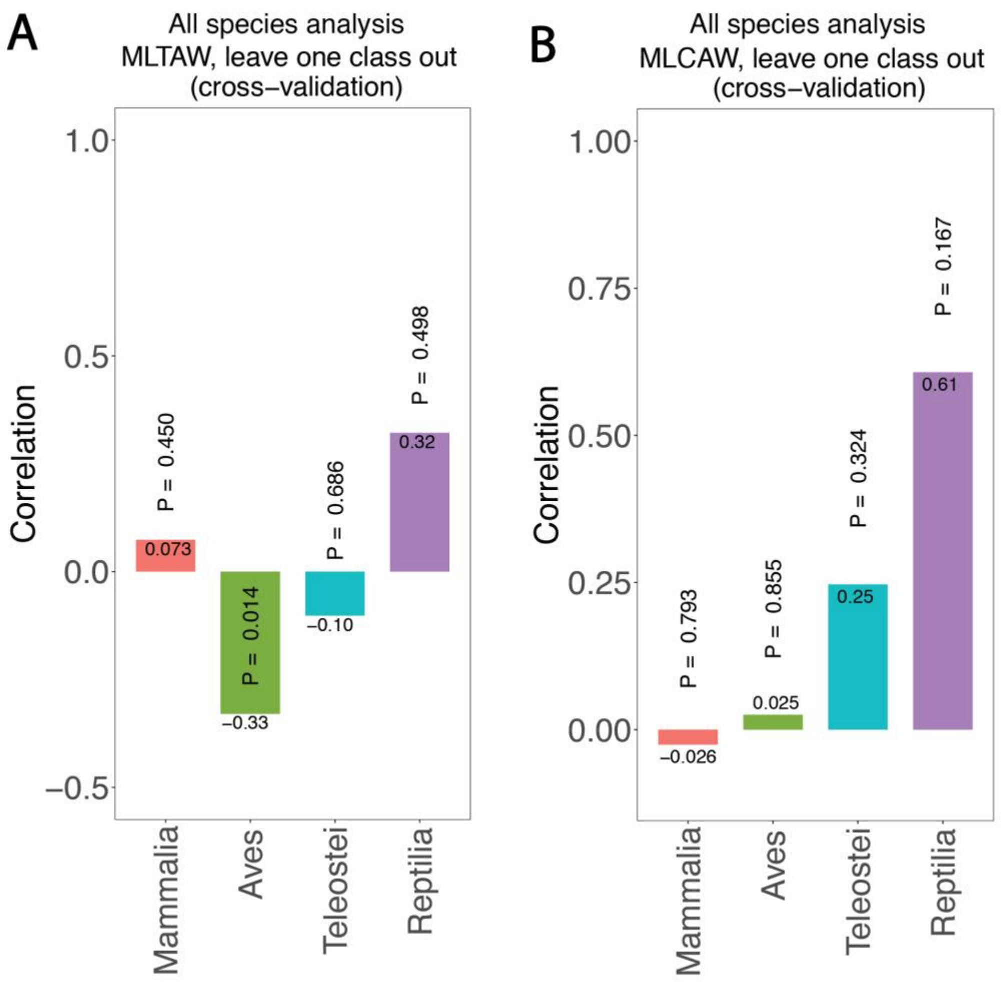


***Figure S10:*** *All species analysis. We predict cancer resistance (CR) scores by identifying PC/NC genes by leaving out one class and testing on that left-out class (cross-validation). We show the accuracies for the following classes: Mammalia (mammals), Aves (birds), Teleostei (fish), Reptilia (reptiles). Spearman's ρ and p-values (P) are reported using the two cancer-resistance estimates:* ***(A)*** *MLTAW; and* ***(B)*** *MLCAW.*

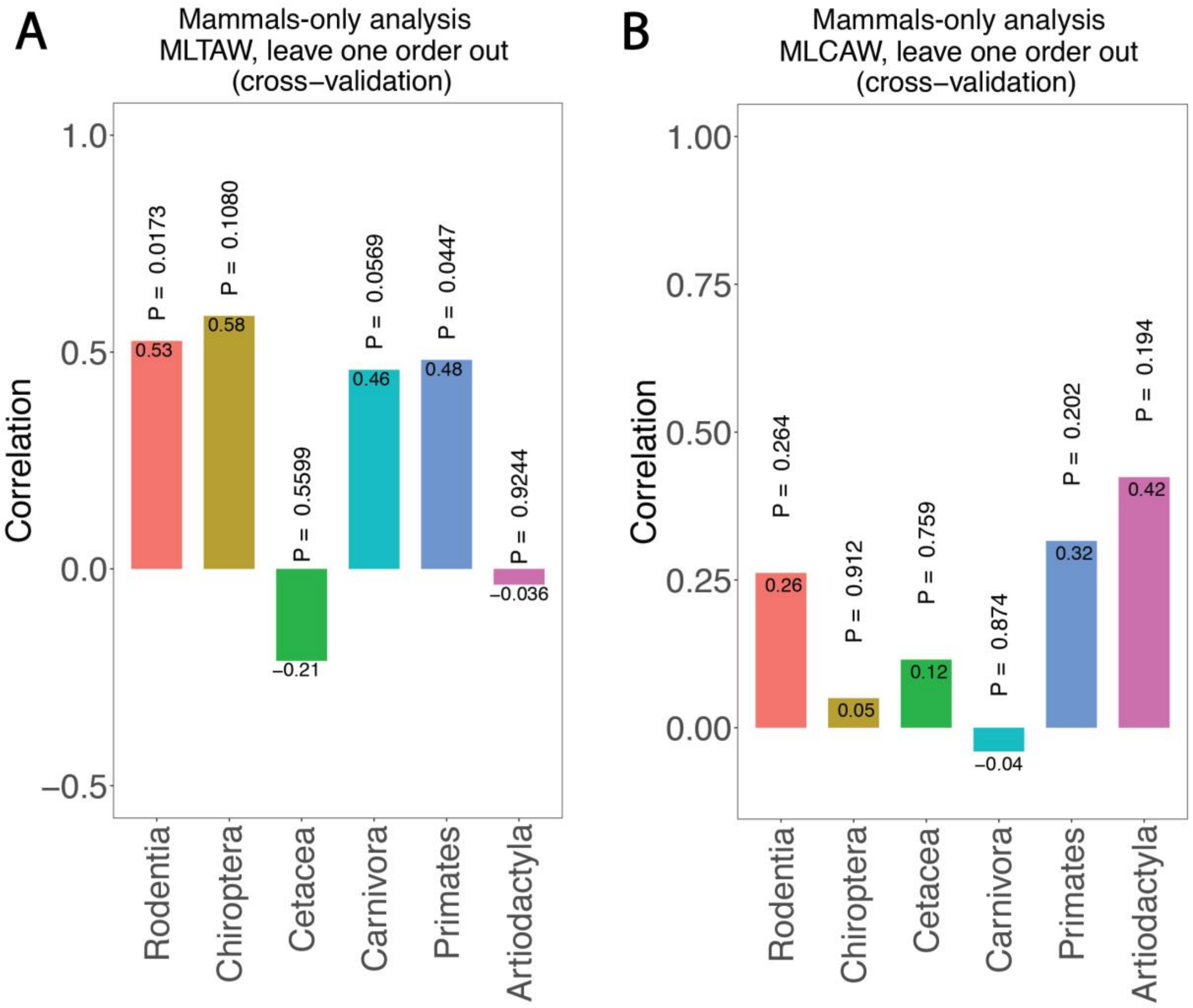


***Figure S11:*** *Mammals-only analysis. We predict cancer resistance (CR) scores by identifying PC/NC genes (using mammalian data) by leaving out one order of mammals and testing on that left-out order (cross-validation). We show the accuracies for the following orders: Rodentia (rodents), Chiroptera (bats), Cetacea (aquatic mammals like whales), Carnivora (carnivores), Primates, Artiodactyla (even-toed hoofed mammals). Spearman's ρ and p-values (P) are reported using the two cancer-resistance estimates:* ***(A)*** *MLTAW; and* ***(B)*** *MLCAW.*

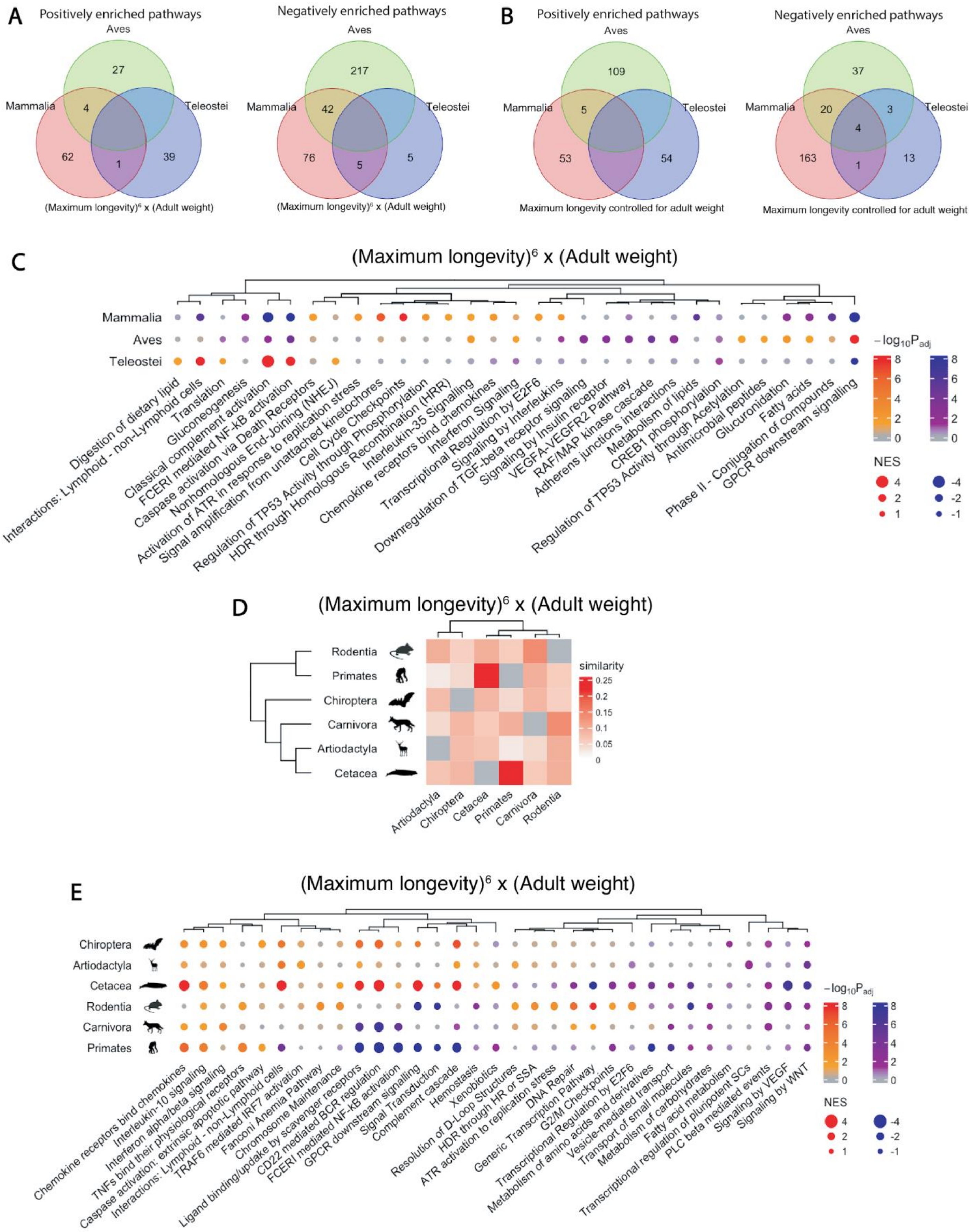
A
Positively enriched pathways
Aves
Mammalia
Teleostei
27
4
62
1
39
(Maximum longevity)⁶ x (Adult weight)
Negatively enriched pathways
Aves
Mammalia
Teleostei
217
42
76
5
5
(Maximum longevity)⁶ x (Adult weight)
B
Positively enriched pathways
Aves
Mammalia
Teleostei
109
5
53
54
Maximum longevity controlled for adult weight
Negatively enriched pathways
Aves
Mammalia
Teleostei
37
20
3
4
163
1
13
Maximum longevity controlled for adult weight
C
(Maximum longevity)⁶ x (Adult weight)
Mammalia
Aves
Teleostei
Digestion of dietary lipid
Interactions: Lymphoid - non-Lymphoid cells
Translation
Gluconeogenesis
Classical complement activation
FCERI mediated NF-kB activation
Caspase activation via Death Receptors
Nonhomologous End-Joining (NHEJ)
Activation of ATR in response to replication stress
Signal amplification from unattached kinetochores
Cell Cycle Checkpoints
Regulation of TP53 Activity through Phosphorylation
HDR through Homologous Recombination (HRR)
Interleukin-35 Signalling
Chemokine receptors bind chemokines
Interferon Signaling
Transcriptional Regulation by E2F6
Signaling by Interleukins
Downregulation of TGF-beta receptor signaling
Signaling by Insulin receptor
VEGFA-VEGFR2 Pathway
RAF/MAP kinase cascade
Adherens junctions interactions
Metabolism of lipids
CREB1 phosphorylation
Regulation of TP53 Activity through Acetylation
Antimicrobial peptides
Glucuronidation
Phase II - Conjugation of compounds
Fatty acids
GPCR downstream signalling
−log10Padj
8
6
4
2
0
NES
4
2
1
-4
-2
-1
D
(Maximum longevity)⁶ x (Adult weight)
Rodentia
Primates
Chiroptera
Carnivora
Artiodactyla
Cetacea
similarity
0.25
0.2
0.15
0.1
0.05
0
Artiodactyla
Chiroptera
Cetacea
Primates
Carnivora
Rodentia
E
(Maximum longevity)⁶ x (Adult weight)
Chiroptera
Artiodactyla
Cetacea
Rodentia
Carnivora
Primates
Chemokine receptors bind chemokines
Interleukin-10 signaling
Interferon alpha/beta signaling
TNFs bind their physiological receptors
Caspase activation: extrinsic apoptotic pathway
Interactions: Lymphoid - non-Lymphoid cells
TRAF6 mediated IRF7 activation
Fanconi Anemia Pathway
Chromosome Maintenance
Ligand binding/uptake by scavenger receptors
CD22 mediated BCR regulation
FCERI mediated NF-kB activation
GPCR downstream signalling
Signal Transduction
Complement cascade
Hemostasis
Xenobiotics
Resolution of D-Loop Structures
HDR through HR or SSA
ATR activation to replication stress
DNA Repair
Generic Transcription Pathway
G2/M Checkpoints
Transcriptional Regulation by E2F6
Metabolism of amino acids and derivatives
Vesicle-mediated transport
Transport of small molecules
Metabolism of carbohydrates
Fatty acid metabolism
Transcriptional regulation of pluripotent SCs
PLC beta mediated events
Signaling by VEGF
Signaling by WNT
−log10Padj
8
6
4
2
0
NES
4
2
1
-4
-2
-1

***Figure S12:*** *Gene set enrichment analysis (GSEA) of the correlation between the gene conservation scores and cancer-resistance estimates including (maximum longevity)$^6$ x (adult weight) (MLTAW), or the residue of maximum longevity after regressing out adult weight (MLCAW), for three classes of species: Mammalia (mammals), Aves (birds), and Teleostei (fish).* ***(A,B)*** *Venn diagram showing the number of positively and negatively enriched gene sets in the three classes based on correlations with: (A) MLTAW and (B) MLCAW.* ***(C)*** *A summary visualization of the GSEA result for the top significantly enriched gene sets in the three classes (Mammalia, Aves, Teleostei) based on correlations with MLCAW. A selected subset of top gene sets are shown to save space, all with adjusted P<0.1 in at least one of the classes. GSEA significance (negative log10 adjusted P values) is encoded by dot color, with two sets of colors (red-orange and blue-purple) representing positive or negative enrichment, respectively; grey color means adjusted P>=0.1. Dot size represents the absolute value of normalized enrichment scores (NES) measuring the effect size of enrichment. The complete GSEA results are given in Table S3.* ***(D)*** *GSEA analysis of the correlation between the gene conservation scores and cancer-resistance estimates such as MLTAW were performed for different orders of mammalian species including Rodentia (rodents), Primates (primates), Chiroptera (bats), Carnivora (carnivores), Artiodactyla (even-toed hoofed animals), and Cetacea (whales). A heatmap showing the similarity (Jaccard index) between the significantly enriched gene sets (FDR<0.1) from each pair of mammalian orders, based on the MLTAW correlation. The dendrogram on the left is the phylogenetic tree of the mammalian orders, and the rows of the heatmap are arranged accordingly. The dendrogram on the top represents the hierarchical clustering of the orders based on their similarities in the GSEA results.* ***(E)*** *A summary visualization of the GSEA result for the top significantly enriched gene sets in the mammalian orders based on MLTAW correlation. A selected subset of top gene sets are shown to save space, all with adjusted P<0.1 in at least one of the orders (complete results in Table S5). The color code and dot size is as described in (C).*

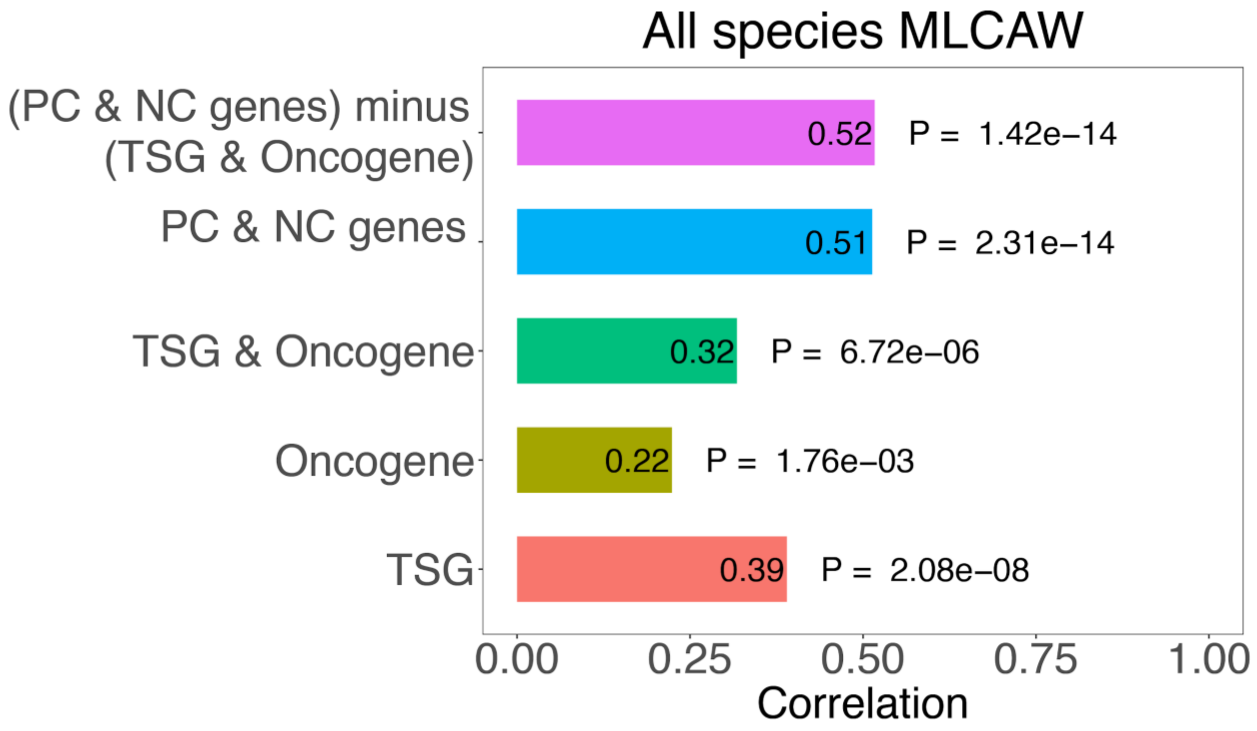


***Figure S13:*** *Spearman's correlation (ρ) in predicting cancer resistance (MLCAW) in all species using only TSGs, only oncogenes, both TSGs and oncogenes, using PC and NC genes in cross validation, using PC and NC genes after removing TSGs and oncogenes in cross validation is shown. Analysis is done using all species.*

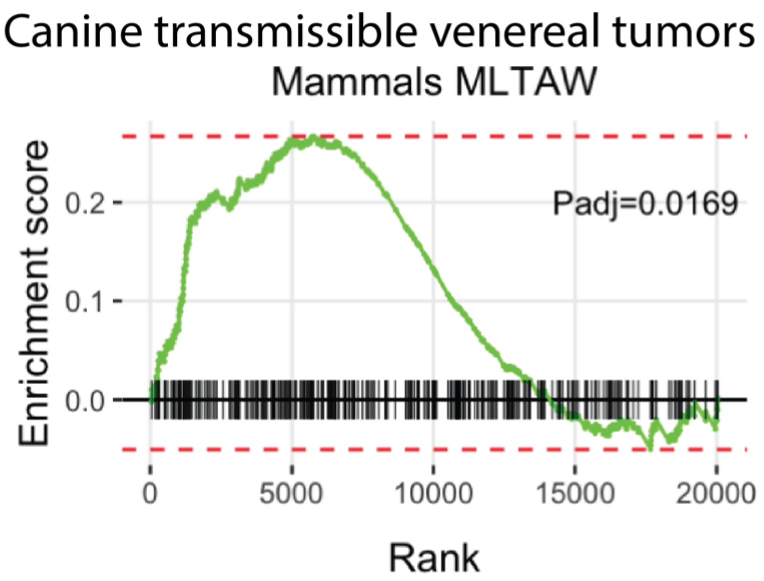


***Figure S14:*** *Gene set enrichment analysis (GSEA) plot showing a significant enrichment of the PC genes in mammals (using MLTAW measure) for the loss-of-function genes observed in canine transmissible venereal tumors.*

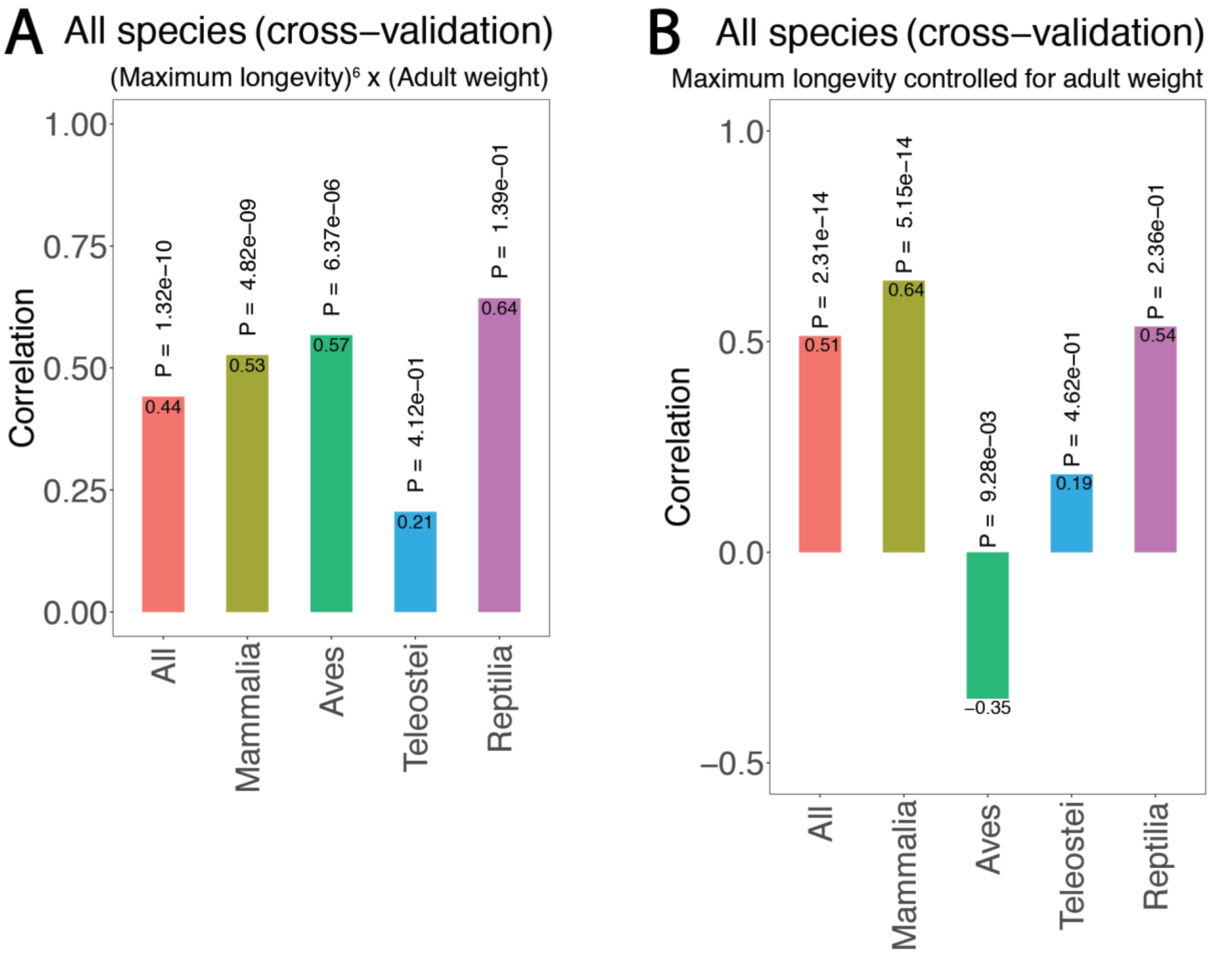


***Figure S15:*** *The cancer resistance (CR) scores predicted on all species (in leave-one-out cross-validation) analysis are individually tested on different classes of species: Mammalia (mammals), Aves (birds), Teleostei (fish), Reptilia (reptiles). Spearman's ρ and p-values (P) are reported using the two cancer-resistance estimates:* ***(A)*** *MLTAW or '(Maximum longevity)$^6$ x (adult weight)'; and* ***(B)*** *MLCAW or 'Maximum longevity controlled for adult weight'.*
